# MONN: a Multi-Objective Neural Network for Predicting Pairwise Non-Covalent Interactions and Binding Affinities between Compounds and Proteins

**DOI:** 10.1101/2019.12.30.891515

**Authors:** Shuya Li, Fangping Wan, Hantao Shu, Tao Jiang, Dan Zhao, Jianyang Zeng

**Affiliations:** Institute for Interdisciplinary Information Sciences, Tsinghua University, Beijing, China; Department of Computer Science and Engineering, University of California, Riverside, CA; Bioinformatics Division, BNRIST/Department of Computer Science and Technology, Tsinghua University, Beijing, China; MOE Key Laboratory of Bioinformatics, Tsinghua University, Beijing, China

## Abstract

Computational approaches for inferring the mechanisms of compound-protein interactions (CPIs) can greatly facilitate drug development. Recently, although a number of deep learning based methods have been proposed to predict binding affinities and attempt to capture local interaction sites in compounds and proteins through neural attentions, they still lack a systematic evaluation on the interpretability of the identified local features. In addition, in these previous approaches, the exact matchings between interaction sites from compounds and proteins, which are generally important for understanding drug mechanisms of action, still remain unknown. Here, we compiled the first benchmark dataset containing the inter-molecular non-covalent interactions for more than 10,000 compound-protein pairs, and used it to systematically evaluate the interpretability of neural attentions in existing prediction models. We developed a multi-objective neural network, called MONN, to predict both non-covalent interactions and binding affinity for a given compound-protein pair. MONN uses convolution neural networks on molecular graphs of compounds and primary sequences of proteins to effectively capture the intrinsic features from both inputs, and also takes advantage of the predicted non-covalent interactions to further boost the accuracy of binding affinity prediction. Comprehensive evaluation demonstrated that while the previous neural attention based approaches fail to exhibit satisfactory interpretability results without extra supervision, MONN can successfully predict non-covalent interactions on our benchmark dataset as well as another independent dataset derived from the Protein Data Bank (PDB). Moreover, MONN can outperform other state-of-the-art methods in predicting compound-protein binding affinities. In addition, the pairwise interactions predicted by MONN displayed compatible and accordant patterns in chemical properties, which provided another evidence to support the strong predictive power of MONN. These results suggested that MONN can offer a powerful tool in predicting binding affinities of compound-protein pairs and also provide useful insights into understanding the molecular mechanisms of compound-protein interactions, which thus can greatly advance the drug discovery process. The source code of the MONN model and the dataset creation process can be downloaded from https://github.com/lishuya17/MONN.

## 1 Introduction

Elucidating the mechanisms of compound-protein interactions (CPIs) plays an essential role in drug discovery and development [1, 2]. Although various experimental assays [3] have been widely applied for drug candidate screening and property characterization, identifying hit compounds from a large-scale chemical space is often time- and resource-consuming. To relieve this bottleneck, computational methods are typically used to reduce time and experimental efforts [4]. For example, it has been shown that effective high-throughput virtual screening can greatly accelerate the lead discovery process [5].

Apart from the binding and functional assays, structure determination of compound-protein complexes can shed light on the molecular mechanisms of CPIs and thus significantly promote the lead optimization process. In particular, based on the molecular basis of CPIs revealed by the complex structures, drug developers can gain better insights into understanding how to improve the design of candidate compounds, for the purpose of enhancing binding specificities or avoiding side effects [6]. However, determining the atomic resolution structures of protein-ligand complexes through currently available experimental techniques, such as X-ray crystallography [7], nuclear magnetic resonance (NMR) [8] and cryo-electron microscopy (cryo-EM) [9], is still time-consuming in practice, resulting in only a limited number of solved structures [10]. Therefore, a natural question arises: can computational virtual screening methods also provide useful mechanistic insights about CPIs in addition to predicting their binding affinities?

Molecular docking (e.g., AutoDock Vina [11] and GOLD [12]) and molecular dynamics (MD) simulations [13] have been popularly used in virtual screening of compound-protein interactions [14]. These methods have inherently good interpretability, as they can predict potential binding poses as well as binding affinities. Despite a number of successful stories about the applications of these structure-based computational methods, they still suffer from several limitations. One major limitation lies in their heavy dependence on the available 3D-structure data of the protein targets. In addition, these molecular docking and MD simulation based methods generally require tremendous computational resources.

To overcome the current limitations of the structure-based computational methods, a number of structure-free models [15–21] have been developed for predicting CPIs. An example is the similarity-based methods that take similarity matrices as descriptors of both compounds and proteins [15, 16]. These methods mainly focus on the global similarities of entire compounds or proteins, while ignoring the detailed compositions of each molecule. Conversely, deep learning based methods [17–19, 21] fully exploit the local features of input compound structures and protein sequences to predict their binding affinities.

A fraction of these structure-free methods make use of neural attentions, which have been widely used in the deep learning community to guide models to focus on those “important” features, and thus increase the interpretability of their prediction results [22, 23]. For the CPI prediction tasks [17–19], attentions are expected to be able to capture the local binding sites mediated by non-covalent interactions (e.g., hydrogen bonds and hydrophobic effects) between compounds and proteins. Although these methods demonstrated that real binding sites of compounds or proteins were enriched in their attention-highlighted regions in a few examples, systematic comparison and evaluation on this learning capacity are still lacked, probably due to the absence of benchmark datasets and evaluation standards. In this paper, we constructed the first benchmark dataset containing pairwise non-covalent interactions between atoms of compounds and residues of proteins from more than 10,000 compound-protein pairs, and comprehensively evaluated the interpretability of different neural attention based frameworks. Interestingly, tests on our constructed benchmark dataset showed that current neural attention based approaches have difficulty in automatically capturing the accurate local non-covalent interactions between compounds and proteins without extra supervised guidance.

Based on this observation, we proposed MONN, a Multi-Objective Neural Network, to learn both pairwise non-covalent interactions and binding affinities between compounds and proteins. MONN is a structure-free model that takes only graph representations of compounds and primary sequences of proteins as input, with capacity to handle large-scale datasets with relatively low computational complexity. The input information is processed by graph convolution networks and convolution neural networks (CNNs), but different from previous CPI prediction methods in the following aspects: 1) MONN uses a graph warp module [24] in addition to a traditional graph convolution module [25] to learn both a global feature for the whole compound and local features for each atom of the compound to better capture the molecular features of compounds; 2) MONN contains a pairwise interaction prediction module, which can capture the pairwise non-covalent interactions between atoms of a compound and residues of a protein with extra supervision from the labels extracted from available high-quality 3D compound-protein complex structures; and 3) in MONN, pairwise non-covalent interaction prediction result is further utilized to benefit the prediction of binding affinities, by effectively incorporating the shared information between compound features and protein features into the downstream affinity prediction module.

Comprehensive cross-validation tests on our constructed benchmark dataset demonstrated that MONN can successfully learn the pairwise non-covalent interactions derived from high-quality structure data, even using the 3D structure-free information as input. We also used an additional test dataset constructed from the PDB to further validate the generalization ability of MONN. Moreover, extensive tests showed that MONN can achieve superior performance in predicting CPI binding affinities, over other state-of-the-art structure-free models. In addition, although the chemical rules, such as the correlation of hydrophobicity scores between compounds and proteins and the preference of atom and residue types for hydrogen bonds and *π*-stacking interactions, are not explicitly incorporated into the prediction framework, such features can still be effectively captured by MONN.

In summary, the following contributions are made in this paper:

1. We combined the predictions of pairwise non-covalent interactions and binding affinities between compounds and proteins into a unified machine learning problem, constructed the first large-scale benchmark dataset for this problem and systematically evaluated the interpretability of neural attentions on this dataset.
2. We developed MONN, a novel deep learning framework to effectively extract molecular features from 3D-structure independent representations of compounds and proteins, and predict both the local atom-level interactions (*i.e.*, pairwise non-covalent interactions) and the global binding strengths (*i.e.*, affinities).
3. The comprehensive tests on the constructed benchmark dataset and validation datasets demonstrated that MONN can outperform other state-of-the-art models in predicting binding affinities, and also accurately capture the pairwise non-covalent interactions between compounds and proteins. These test results suggested that MONN can provide a useful tool for effectively modeling CPIs both locally and globally, and thus greatly facilitate the drug discovery process.

## 2 Methods

### 2.1 Problem formulation

MONN is an end-to-end neural network model with two training objectives. One objective is to predict the non-covalent interactions between the atoms of a compound and the residues of its protein partner. To describe the non-covalent interactions in a computational manner, we define a pairwise interaction matrix: for a compound with *N*_*a*_ atoms, and a protein with *N*_*r*_ residues, their pairwise interaction matrix ***P*** is defined as an *N*_*a*_ × *N*_*r*_ binary matrix, in which each element *P*_*ij*_ (*i* = 1, 2, …, *N*_*a*_ and *j* = 1, 2, …, *N*_*r*_) indicates whether there exists a non-covalent interaction (1 for existence, 0 otherwise) between the *i*-th atom of the compound and the *j*-th residue of the protein when forming a complex structure. The other objective is to predict binding affinities (e.g., K_i_, K_d_ or IC_50_), which can also be regarded as a global measurement of the binding strength, between a protein and its ligand. Binding affinity can be denoted by a real number *a* ∈ ℝ.

A chemical compound with *N*_*a*_ atoms can be represented by a graph *G* = {*V, E*}, where each node *v*_*i*_ ∈ *V* (*i* = 1, 2, …, *N*_*a*_), corresponds to the *i*-th atom in the compound, and each edge 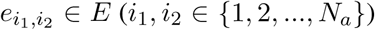 corresponds to a chemical bond between the *i*_1_-th and the *i*_2_-th atoms. A protein with *N*_*r*_ residues can be represented by a string of its primary sequence, denoted by 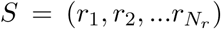, where each *r*_*j*_ (*j* = 1, 2, …, *N*_*r*_) is either one of the 20 standard amino acids, or a letter ‘X’ for any non-standard amino acid. Given a graph representation of a compound and a string representation of a protein sequence, our model is expected to output a predicted pairwise non-covalent interaction matrix 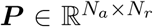 and an estimated affinity value *a* ∈ ℝ.

### 2.2 The network architecture of MONN

Our model consists of four modules (Figs 1, S1-S4): i) a graph convolution module for extracting the features of both individual atoms and the whole compound from a given molecular graph, ii) a convolutional neural network (CNN) module for extracting the features of individual residues from a given protein sequence, iii) a pairwise interaction prediction module for predicting the probability of the non-covalent interaction between any atom-residue pair from the previously learned atom and residue features, and iv) an affinity prediction module for predicting the binding affinity between the given pair of compound and protein, using the previously extracted molecular features, as well as the derived pairwise interaction matrix.

**Fig. 1.**
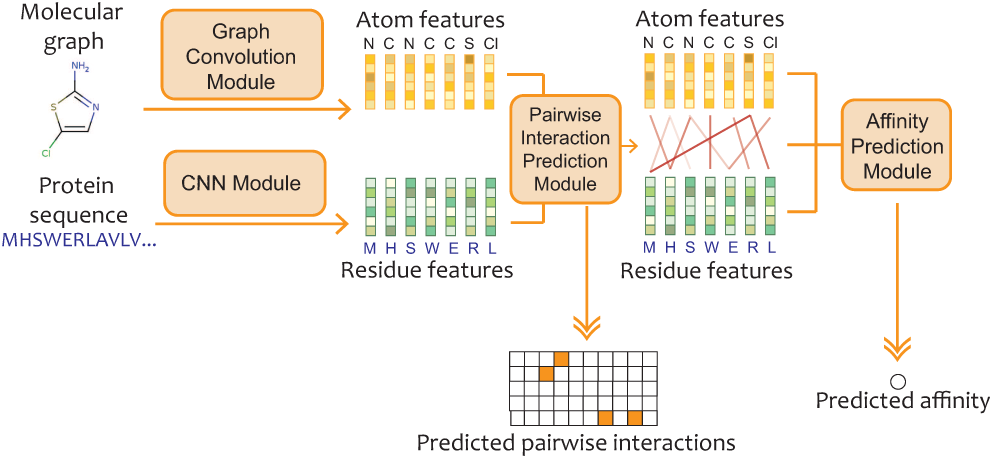
The architecture overview of MONN. Given a compound-protein pair, a graph convolution module and a convolution neural network (CNN) module are first used to extract the atom and residue features from the input molecular graph and protein sequence, respectively. Then, these extracted atom and residue features are processed by a pairwise interaction prediction module to derive the predicted pairwise interaction matrix, which enables one to construct the links between atoms of the compound and residues of the protein. Finally, an affinity prediction module is used to integrate information from atom features, residue features and the previously derived pairwise interactions to obtain the predicted binding affinity.

### The graph convolution module

The graph convolution module (Fig S1) takes the graph representation *G* = {*V, E*} of a compound as input. More specifically, each node (*i.e.*, atom) *v*_*i*_ ∈ *V* is initially represented by a feature vector 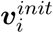 of length 82, which is the concatenation of one-hot encodings representing the atom type, degree, explicit valence, implicit valence and aromaticity of the corresponding atom. Then, the initial atom features are transformed into 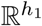 (*h*_1_ is the hidden size) by a single-layer neural network:

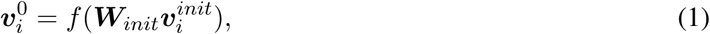

where *f* (·) stands for the leaky ReLU activation function *f* (*x*) = max(0, *x*) + 0.1 min(0, *x*), and 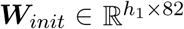. Note that for all the single-layer neural networks in this paper, unless otherwise stated, *f* (·) stands for the leaky ReLU activation function, ***W***_*x*_ (*x* can be any subscript) stands for the learnable weight parameters, and the bias terms are omitted for clarity.

Each edge (i.e., chemical bond) 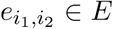 is represented by a feature vector 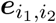 of length 6, which is the concatenation of one-hot encodings representing the bond type (single, double, triple or aromatic) and other properties, e.g., whether the bond is conjugated and whether it is in a ring.

The atom features are then processed by *L* iterations of graph convolution to produce a set of updated atom features 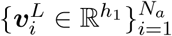 and a super node feature 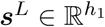, which is an overall feature representation for the compound of interest. Note that the bond features are not updated during the whole process.

At each iteration of graph convolution, the atom features are sequentially updated using both a basic message passing unit [25] and a graph warp unit [24]. The message passing unit executes the following two steps to extract the local features from the given graph: gathering information and updating information. During the first step (*i.e.*, gathering information), each atom *v*_*i*_ gathers local information 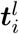 from both its neighboring atoms and bonds, that is,

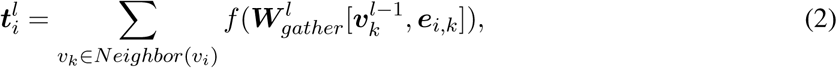

where *i* = 1, 2, …, *N*_*a*_, *l* = 1, 2, …, *L*, 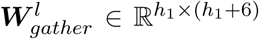, *Neighbor*(*v*_*i*_) stands for the set of neigh boring atoms of 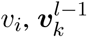 represents the feature of atom *v*_*k*_ from the (*l* − 1)-th layer, and [·, ·] stands for the concatenation operation. In the second step (*i.e.*, updating information), the gathered information and the atom features learned from the previous iteration are then processed to obtain the updated features 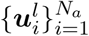 at each iteration *l*, that is,

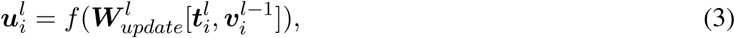

where *i* = 1, 2, …, *N*_*a*_, *l* = 1, 2, …, *L*, and 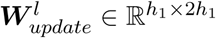.

The graph warp unit [24] further improves the performance (the results of the corresponding ablation studies are shown in Figs S5-S6) of graph convolution networks by introducing a super node *s*, which captures the global feature for the compound of interest. Through information sharing between the super node and all the atoms, distant atoms in the graph can communicate effectively and efficiently through this super node, and thus a global feature can be extracted based on this technique [24]. More specifically, this information sharing operation takes the atom features 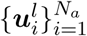 updated in current iteration *l* and the super node feature *s*^*l*−1^ from the previous iteration *l* − 1 as input, and outputs both updated atom features 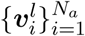 and super node feature *s*^*l*^. Details about the implementation of this graph warp unit can be found in Supplementary Note S1.1.

After *L* iterations of graph convolution as described above, the final atom features 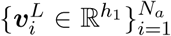 and the super node feature 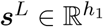 are generated and then fed into the downstream modules. In the remaining part of this paper, we will drop the superscript *L* for clarity.

### The CNN module

The protein sequence is first encoded using the BLOSUM62 matrix [26], that is, the initial feature of each residue is represented by the corresponding column of the BLOSUM62 matrix. The features of non-standard amino acids are zero-initialized. We use this encoding strategy instead of the commonly used one-hot encoding scheme for protein sequences, mainly because the BLOSUM62 matrix is a 20 × 20 matrix that has encoded the evolutionary relationships between amino acids, while the one-hot encoding scheme lacks such information. Then, the initial features are updated through typical 1-D convolution layers [27] with a leaky ReLU activation function. The specific architecture of the employed convolution neural network is determined by three hyper-parameters, including the number of convolution layers, the number and the size of filters in each layer (Supplementary Notes S2.2). In the end, we obtain the final output features 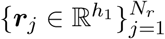 for all the residues along the protein sequence (Fig S2).

### The pairwise interaction prediction module

To predict the pairwise interactions between a given compound-protein pair, the pairwise interaction prediction module (Fig S3) uses the atom features 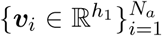 and the residue features 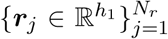 derived from the modules described above. The atom and residue features are first transformed into a compatible space by two single-layer neural networks separately. Then, the predicted probability of the interaction between an atom *v*_*i*_ and a residue *r*_*j*_ is derived based on the inner product between the transformed atom and residue features, normalized by a sigmoid function:

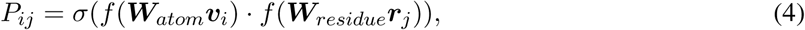

where *i* = 1, 2, …, *N*_*a*_, *j* = 1, 2, …, *N*_*r*_, 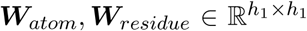, *σ*(·) represents the sigmoid function 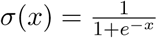, and · denotes the inner product.

### The affinity prediction module

The affinity prediction module (Fig S4) integrates information from not only the previously learned atom features 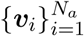, the super node feature ***s*** and the residue features 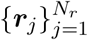, but also the predicted pairwise interaction matrix ***P***. Intuitively, ***P*** can be used to construct the links and share information between atom and residue features, which may thus provide additional useful information for predicting the binding affinity.

First, the atom features 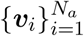 and the super node feature ***s***, as well as the residue features 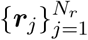, are transformed into a compatible space for affinity prediction by single-layer neural networks:

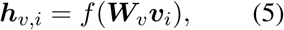

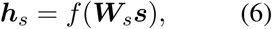

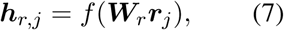

where 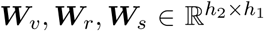, and *h*_2_ is the size of hidden units in the single-layer neural networks used in the affinity prediction module.

Next, we generate a fixed-size feature representation for each compound and protein, from a list of transformed atom and residue features, using an attention mechanism that has been widely used to enhance the performance of deep learning. In particular, the neural attention mechanism is introduced to weigh the contributions of features from individual atoms and residues, which has been proved to be more effective than simply averaging all the atom and residue features (the results of the corresponding ablation studies are shown in Figs S5-S6). The dual attention network (DAN) [28] is a recently published method that can produce attentions for two given related entities (each with a list of features). For example, given an image with a sentence annotation, DAN generates a textual attention for the word features of the sentence and a visual attention for the spatial features of the image. Here, we modified the DAN framework by utilizing the predicted pairwise interaction matrix to construct the direct links between atoms and residues. Information passing is thus enabled by gathering features of interaction partners through such links for each atom of the compound and each residue of the protein. The passed information is then incorporated into the calculation of compound and protein attentions by DAN (Details can be found in Supplementary Note S1.2.).

Through the compound and protein attentions, the atom features and the residue features can be reduced into the fixed-size representations of the input compound graph (denoted by ***h***_*c*_) and protein sequence (denoted by ***h***_*p*_):

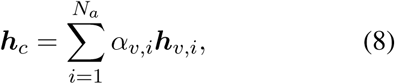

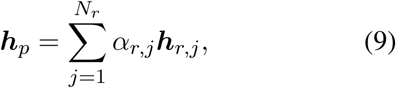

where 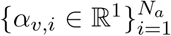 and 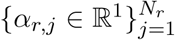 are compound and protein attentions generated by the modified DAN.

Then, ***h***_*c*_ is concatenated with the transformed super node feature ***h***_*s*_ to obtain a combined representation of the compound features (*i.e.*, [***h***_*c*_, ***h***_*s*_]). To explore the relationship between this combined representation of the compound features and the representation of the protein features, we calculate their outer product, normalized by a leaky ReLU activation function *f*, and then followed by a linear regression layer to predict the binding affinity, that is,

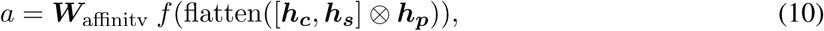

where ⊗ denotes the outer product, flatten(·) reshapes the result of the outer product into a column vector of length 2*h*_2_^2^, and 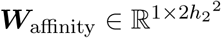.

### 2.3 Training

For a training dataset with *N* samples (*i.e.*, compound-protein pairs), we minimize the cross-entropy loss for pairwise non-covalent interaction prediction, which is defined as

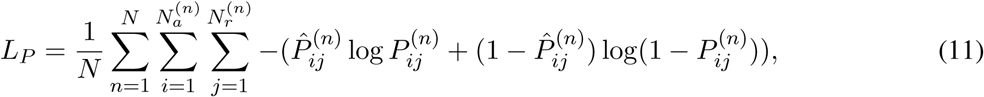

where 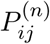 and 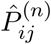 stand for the predicted probability and the true binary label of the interaction between the *i*-th atom and the *j*-th residue in the *n*-th sample, respectively, and 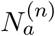 and 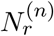 stand for the total number of atoms in the compound and the total number of residues in the protein in the *n*-th sample, respectively.

For binding affinity prediction, the objective is to minimize the mean squared error, which is defined as

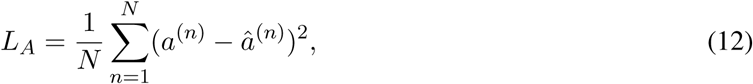

where *a*^(*n*)^ and *â*^(*n*)^ stand for the predicted affinity and the true affinity label for the *n*-th sample, respectively.

In a multi-objective training process, we aim to minimize the combination of two losses to further enhance the binding affinity prediction, that is,

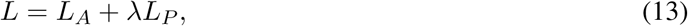

where *λ* stands for a weight parameter controlling the contribution of *L*_*p*_ to the final affinity prediction. During the training process, we used a mini-batch stochastic gradient descent scheme to optimize the model parameters. A single MONN model can be trained within an hour on a Linux server with 48 logical CPU cores and one Nvidia Geforce GTX 1080Ti GPU. More details about training and hyper-parameter calibration can be found in Supplementary Note S2.

### 2.4 Construction of the benchmark dataset

Our benchmark dataset was constructed based on PDBbind v2018 [29, 30], which provides a high-quality set of protein-ligand complexes with available structure data and corresponding binding affinities. For complexes in the PDBbind dataset, we obtained their 3D structures from the RCSB PDB [10] and then extracted the non-covalent interactions between compounds and proteins using PLIP [31]. After considering the atoms types, distances and bond angles, PLIP recognized seven types of non-covalent interactions, *i.e.*, hydrogen bond, hydrophobic interaction, *π*-stacking, *π*-cation, salt bridge, water bridge and halogen bond. For the identified atoms from the compounds and residues from the proteins which are involved in the non-covalent interactions, we mapped their indices into the graph representations of compounds and protein sequences derived from UniProt [32] to construct the corresponding pairwise non-covalent interaction labels. In total we obtained 13,306 compound-protein pairs with binding affinity values, and 12,738 of them had corresponding pairwise interaction labels. More details about the dataset construction process can be found in Supplementary Note S3 and Fig S7.

## 3 Results

### 3.1 Systematic evaluation of the interpretability of neural attentions in CPI prediction models

A number of deep learning-based methods [17–21] have been developed previously for modelling compound-protein interactions from 3D structure-free inputs. Despite their success in predicting binding affinities with relatively low computational complexity, interpretability is still considered as a challenge for these structure-independent methods. Several recent studies [17–19] sought interpretability by incorporating neural attentions (*i.e.*, weighing the contributions of individual elements in the given input to the final predictions) into their model architectures. For example, Tsubaki et al. [17] developed an end-to-end neural network with attentions for protein sequences, and they showed two examples in which their attention highlighted regions were able to capture the real interaction sites in proteins. The method developed by Gao et al. [18] involves both compound and protein attentions, and by visualizing the attention weights, the authors demonstrated that their derived attention highlighted regions can successfully identify the interaction interface in a compound-protein complex. DeepAffinity [19] reported an enrichment of true interaction sites in those regions with high attention scores in protein sequences for several examples.

However, there are still some limitations in the previous studies about the interpretability of the deep learning based CPI prediction methods. First, in these studies, the interpretability of neural attentions was evaluated only through one or several examples, and not comprehensively assessed by a large-scale benchmark dataset. In addition, the evaluations were conducted only by visualizing the attention weights [17, 18] or calculating the enrichment scores [19], thus lacking a unified standard for systematically evaluating the interpretability of different attention-based models. More importantly, in these existing studies, attention weights were mainly used to infer the positions of the interaction sites, but the exact matchings between them (*i.e.*, the pairwise interactions, as defined in Methods) still remain unknown.

To overcome these limitations, we conducted a systematic analysis to evaluate the interpretability of the neural attentions. We first constructed a benchmark dataset containing labels of pairwise non-covalent interactions for about 13,000 compound-protein complexes with available atom-resolution structures (also see Methods and Supplementary Notes S3.1). The non-covalent interaction labels of a compound-protein pair include a binary pairwise interaction matrix (as described in Methods), and the interaction sites derived from this pairwise interaction matrix by maximizing over rows or columns (Fig S7). Then the interpretability was evaluated from the following three aspects: the ability of attentions to capture the interaction sites in compounds (at atom level), the interaction sites in proteins (at residue level), and the pairwise interactions between compounds and proteins. For these binary classification problems, we mainly used the average AUC scores (*i.e.*, averaging over all the compound-protein pairs in the test data) for performance evaluation. In addition, as in DeepAffinity [19], we also calculated the enrichment score, which was defined as the fold change of the precision score of the trained model over the expected precision of random predictions (more details on these metrics can be found in Supplementary Notes S4.1).

Four different types of neural attentions used in existing compound-protein interaction prediction models were evaluated, including the method by Tsubaki et al. [17], the method by Gao et al. [18], the separate and joint attentions proposed in DeepAffinity [19]. Details about the implementations of these neural attentions can be found in Supplementary Notes S4.2. The attention weights were obtained after training the models using the binding affinity labels, that is, without extra supervision from the pairwise interaction labels. The clustering-based cross-validation procedure [33] was used during the training process, which ensures that similar compounds (or/and proteins) in the same clusters were not shared between training and test sets. Three cross-validation settings were used in the evaluation, including the new-compound setting, in which the test compounds were never seen in the training process, the new-protein setting, in which the test proteins were never seen in the training data, and the both-new cross-validation setting, in which both compounds and proteins in the test data were never seen during training. More details about the cross-validation procedure can be found in Supplementary Notes S2.1 and Fig S8.

Under different prediction tasks and cross-validation settings, all the four types of neural attentions achieved average AUC and enrichment scores around 0.5 and 1, respectively, which were close to the scores of random predictions (Fig 2 and Figs S9-S14). These results suggested that, although the attention highlighted regions and the real binding sites displayed accordance in some cases [17–19], they only showed poor correlation in our comprehensive test on a large-scale dataset. Thus, it seems not possible to derive the accurate predictions of non-covalent interactions between compounds and proteins from the attention-based models trained using only binding affinities.

**Fig. 2.**
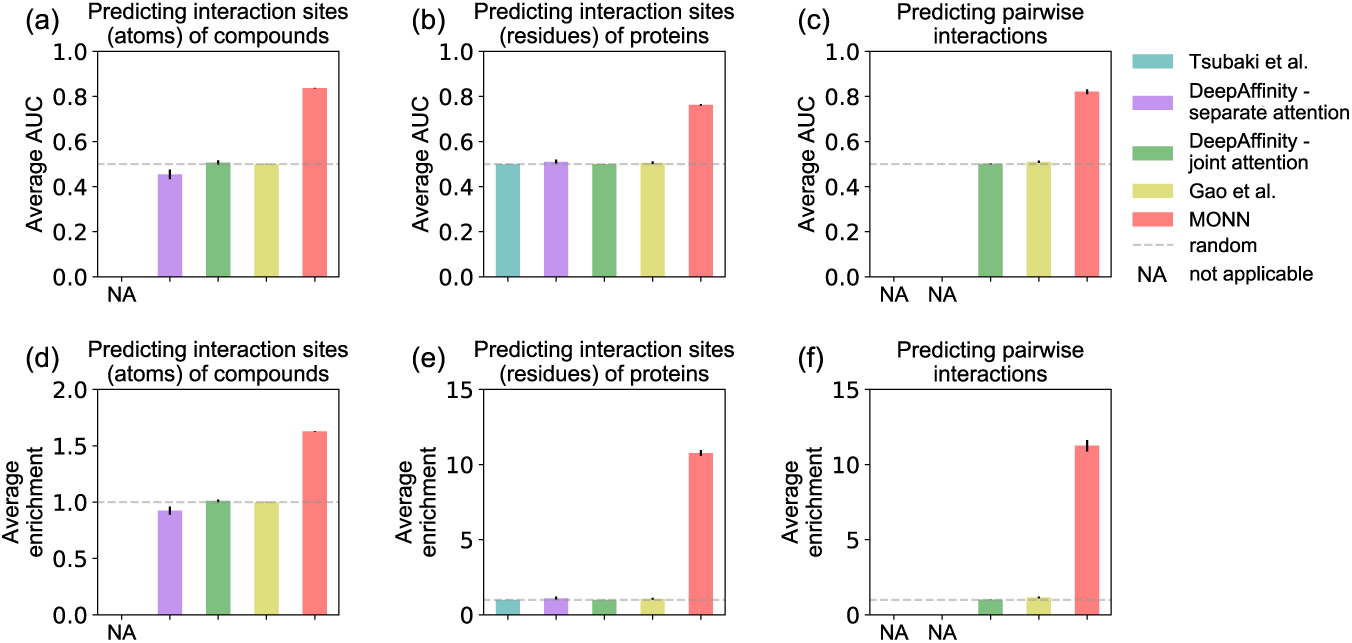
Average AUC scores (a-c) and average enrichment scores (d-e) for evaluating four neural attentions and MONN for the prediction of interaction sites (atoms) in compounds under the new-compound setting (a,d), interaction sites (residues) in proteins under the new-protein setting (b,e), and pairwise non-covalent interactions between compounds and proteins under the both-new setting (c,f). The mean values and standard deviations over 10 repeats of cross-validation with clustering threshold 0.3 are plotted. The ratios of positive and negative labels are about 1:1.44, 1:46.5 and 1:605 under these three cross-validation settings, respectively.

### 3.2 Performance evaluation on pairwise non-covalent interaction prediction by MONN with extra supervision

Based on the above observation that neural attentions cannot automatically capture the non-covalent interactions between compounds and proteins, we speculated that extra supervision information can be used to guide our model to capture such local interactions. Instead of using attention mechanisms, MONN uses an individual module (*i.e.*, the pairwise interaction prediction module) to learn the pairwise non-covalent interactions from given labels (Fig 1, Methods). Meanwhile, through marginalizing the predicted pairwise interaction matrix, the predicted interaction sites in either compounds or proteins can also be derived.

The cross-validation settings and the metrics for evaluating our model were the same as described in the previous section and Supplementary Notes S2.1. As shown in Fig 2, our model achieved average AUC scores of 0.837, 0.763 and 0.821, and average enrichment scores of 1.63, 10.8 and 11.3 under the three application settings, respectively. Note that the values of the enrichment scores were not comparable among these three settings, due to the different ratios of positive-negative labels (Supplementary Notes S4.1). A more comprehensive comparison test (Fig S9-S14) on our model and different neural attentions was performed for different prediction goals, cross-validation settings, and clustering thresholds, which showed that the predictions of MONN are effective and robust (average AUC scores decreased less than 5% with the clustering threshold increasing from 0.3 to 0.6). These results suggested that, while the neural attentions cannot interpret the non-covalent interactions, MONN is able to accurately predict such interactions between compounds and proteins under different cross-validation settings.

To further examine the generalization ability of our model, we also validated MONN on an additional independent dataset containing pairwise non-covalent interactions between compounds and proteins. As our training data (*i.e.*, the PDBbind dataset) included all the high-quality structures of compound-protein complexes originally downloaded from the PDB [10] before 2018, we also constructed an additional test dataset by collecting all the compound-protein complexes from the PDB with the release date from Jan 1st, 2018 to March 31st, 2019 (Supplementary Notes S3.2). In this extra test, MONN achieved average AUC 0.859 and average enrichment score 112.47 for the 1843 in predicting pairwise interactions of compound-protein pairs on this additional dataset (Fig 3a).

**Fig. 3.**
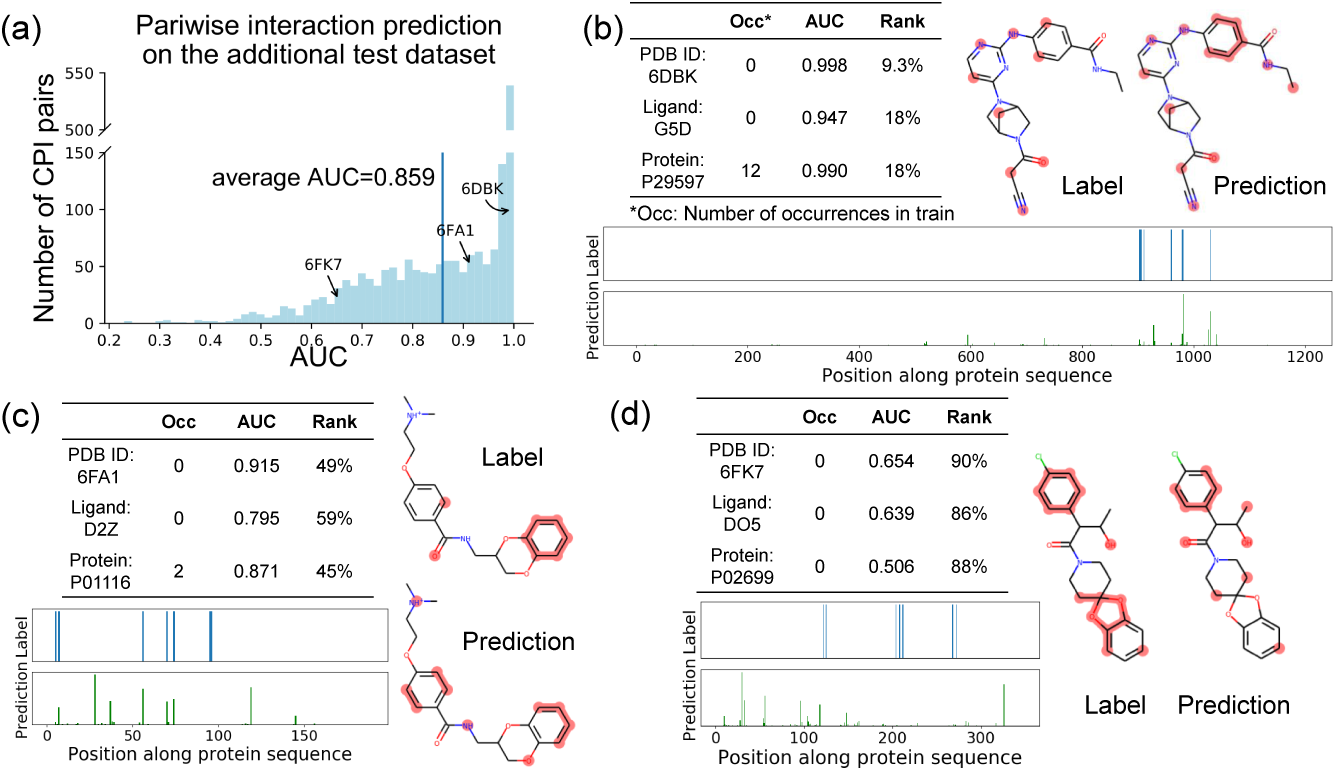
Performance evaluation of MONN on the additional test dataset. (a) The distribution of AUC scores for all the compound-protein pairs. Three example pairs ranked around 10%, 50% and 90% in terms of AUC scores for the pairwise interaction prediction are shown in (b-d), respectively. For each example pair, the numbers of occurrences of the same pair, the compound and the protein in training data are listed, as well as the AUC scores and the corresponding ranks for the predicted pairwise interactions, the interaction sites (atoms) in compounds, and interaction sites (residues) in proteins. In the compound structures, true labels and top 40% predicted interaction sites are marked in red using RDKit [34]. In the protein sequences, the true labels and the MONN predicted scores for individual positions are plotted.

To visualize the prediction results of our model, we selected three representative compound-protein pairs ranked around 10%, 50% and 90% in terms of the AUC scores, and plotted the corresponding true labels and the predicted interaction sites in the compound structures and protein sequences (Fig 3b-d). The example pair ranked around top 10% was a tyrosine kinase inhibitor binding to TYK2 (Fig 3b, PDB ID: 6DBK) [35]. The top 40% of the predicted interaction sites (atoms) in the compound covered all the true interaction sites, and the high prediction scores were also appeared around the true interaction sites along the protein sequence. The example pair ranked around the median prediction score contained a compound binding to KRAS (Fig 3c, PDB ID: 6FA1) [36]. The predicted interaction sites of the compound had several overlaps with true interaction sites (5/8 recall), but also with several false positives. For example, the positively charged group in the compound was predicted as an interaction site, which is actually located outside the binding pocket. The predicted interaction sites (residues) of the protein had several overlaps with the true labels, but also with a number of false positives. The example pair ranked around 90% was a ligand binding to rhodopsin (Fig 3d, PDB ID: 6FK7) [37]. The deviation of the predicted interaction sites from true labels in this example was probably due to the scarcity of training data to support these predictions. These visualization results demonstrated that the accuracies of MONN predictions were consistent with their corresponding rankings in AUC scores. Overall, the above comprehensive validation tests supported the strong predictive power of MONN.

### 3.3 Performance evaluation of binding affinity prediction by MONN with single- and multi-objective learning

In this section, we examined the affinity prediction performance of MONN, and compared with other state-of-the-art models. For the binding affinity prediction task, we separated our PDBbind-derived dataset into two subsets, named IC50 (which contained IC_50_ values) and KIKD (which contained both K_i_ and K_d_ values). The main reason for such a separation was that IC_50_ values are generally dependent on experimental conditions, and thus often considered noisier than the measured K_i_ and K_d_ values. Here, the IC50 dataset with the new-compound setting and clustering threshold 0.3 was used for hyper-parameter calibration. More details about training and hyper-parameter selection can be found in Supplementary Note S2.2.

We considered the following state-of-the-art baseline methods for comparison: the similarity-based kernel method CGKronRLS [15], and the deep learning based methods, including DeepDTA [21], Tsubaki el al.’s method [17] and DeepAffinity [19]. As in the previous sections, MONN and baseline methods were evaluated under three different settings of clustering-based cross-validation (*i.e.*, new-compound, new-protein and both-new), in terms of Pearson correlation (Fig 4) and root mean squared error (RMSE, Fig S15). To investigate whether involving the extra supervision from the pairwise interaction labels can help predict the binding affinities, we mainly tested MONN under two conditions: one was a single objective model, denoted as MONN_single_, which used only the affinity labels as supervision information; the other was a multi-objective model, denoted as MONN_multi_, which considered both pairwise interactions and binding affinities into the training objectives.

**Fig. 4.**
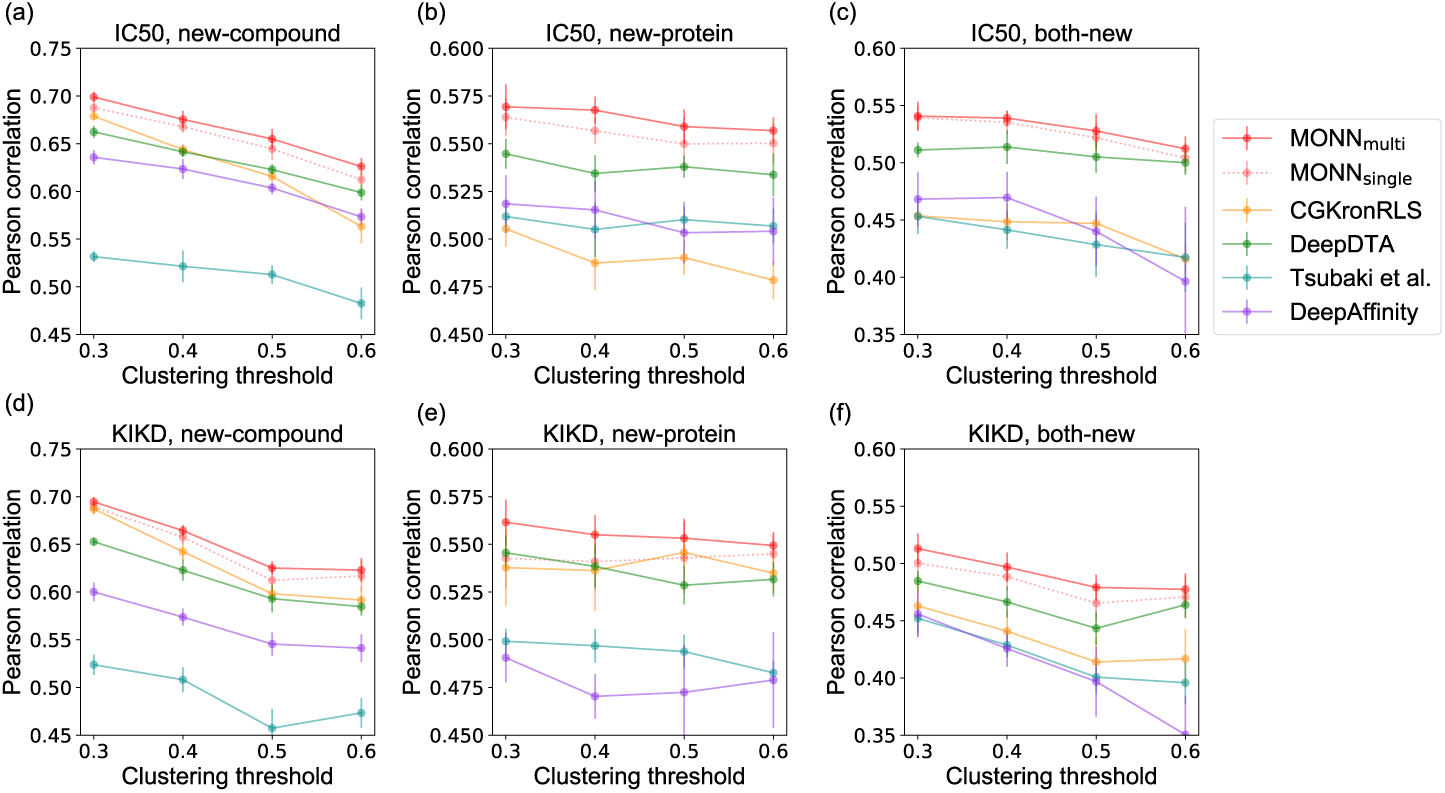
Performance evaluation on binding affinity prediction on the IC50 dataset (a-c) and the KIKD dataset (d-f). Pearson correlations achieved by MONN with single (denoted as MONN_single_) or multiple (denoted as MONN_multi_) training objectives and four baseline methods, under three different cross-validation settings and four different clustering thresholds are shown. The mean values and standard deviations over 10 repeats of cross-validation are plotted.

Our tests showed that both MONN_single_ and MONN_multi_ generally outperformed other baseline methods in all the three cross-validation settings with different clustering thresholds, on both IC50 and KIKD datasets (Fig 4). In particular, compared to the baseline methods, the multi-objective model (MONN_multi_) achieved an increase in Pearson correlation by up to 3.6% (average 2.3%). In addition, the multi-objective model performed slightly better than the single objective one, which indicated that involving extra supervision information from pairwise interaction labels can further improve the binding affinity prediction.

Since compound-protein complexes generally have limited structural availability, we further tested our model on a large-scale structure-free CPI dataset. To our best knowledge, among the baseline methods, only DeepAffinity has been evaluated previously on a large dataset with more than 260,000 training samples and more than 110,000 test samples, with the IC_50_ values derived from the BindingDB database [38]. We followed DeepAffinity’s experimental settings and also tested MONN and DeepDTA on the same dataset. Tsubaki et al.’s method and CGKronRLS are not suitable for this test mainly due to their limited scalability in processing such a large dataset. To make a fair comparison, we also evaluated an ensemble version (*i.e.*, averaging predictions from several single models) of MONN on this BindingDB dataset, as in the DeepAffinity paper [17]. As shown in Table S3, the ensemble version of MONN achieved the best Pearson correlation and RMSE on the BindingDB dataset. This comparison result suggested that, MONN can achieve better performance than the state-of-the-art baseline methods even when the structure data is not available. More details about this test can be found in Supplementary Notes S5.

### 3.4 MONN captures the global molecular property

From the perspective of chemical properties, the size, shape and hydrophobicity of a protein binding pocket are essential for its interaction with a compound [39]. Information about the size and shape of a binding pocket is usually hard to derive only based on raw sequences, so we mainly examined the hydrophobicity of the potential binding residues predicted by MONN, through calculating the correlation between the hydrophobicity scores of the entire compounds and the average hydrophobicity scores of the predicted interaction sites (residues) in the proteins. Here, the hydrophobicity of the compound was measured by the logP value calculated by RDKit [34], which is defined as the log ratio of the solubility of the compound in organic solvent (e.g., 1-octanol) against water [40]. The hydrophobicity of the (predicted) interacting sites of a protein is defined as the average hydrophobicity score over the corresponding side chains [41]. Here, the predicted interaction sites of proteins were selected from the top scored atom-residue pairs in the predicted pairwise interaction matrix ***P***, according to a cut-off value of mean(***P***) + 3 × std(***P***), where std(·) stands for the standard deviation.

The true interaction sites of proteins derived from the solved structures in the benchmark dataset showed a certain level of correlation (Pearson correlation 0.487) in hydrophobicity with their ligands (Fig S16a). As a control, no significant correlation can be observed from randomly chosen residues (Fig S16b). The interaction sites of proteins predicted by MONN had similar correlations in hydrophobicity scores with their ligands (0.515 from cross-validation and 0.499 from the additional test dataset, Fig S16c-d), close to that of true labels.

### 3.5 MONN captures the chemical rules of non-covalent interactions

The rules of non-covalent interactions and the information of interaction types between compounds and proteins are not explicitly incorporated into MONN. Nevertheless, we examined whether MONN can automatically capture such chemical rules. In particular, here, we analyzed the preference of interaction partners for the atoms which can form hydrogen bonds or *π*-stackings.

We first define the conditional likelihood score, which characterizes the preference of residues with a specific property under given atom type of their interaction partners, that is, *p*(residue property=*x*|atom property=*y*) = (Number of residues ∈ *S*(*x*) that interact with the atoms of property *y*)*/*(Total number of residues interacting with the atoms of property *y*), where *S*(*x*) represents the set of residues whose side chains contain at least one kind of elements satisfying the property *x*. Details about the calculation of this conditional likelihood score can be found in Supplementary Notes S6.

A hydrogen bond forms between a hydrogen donor group and an acceptor group. When the atoms from compounds are hydrogen-bond acceptors, the conditional likelihood of hydrogen-bond donor residues as their interaction partners (0.63, calculated using true labels) is much higher than the control residues (0.38, calculated using the randomly chosen residues, Fig S17a). The conditional likelihood scores calculated using MONN predicted interaction sites were also relatively high (0.62 from cross validation and 0.64 from the additional test, Fig S17a). Similarly, hydrogen-bond acceptor residues from the MONN prediction results also had significantly higher conditional likelihood scores than the random control when their interaction partners were the hydrogen-bond donor atoms from the compounds (Fig S17b). For *π*-stacking interactions that occur between aromatic rings, similar conclusions can be drawn: MONN can capture the preference of aromatic residues as interaction partners of aromatic atoms (detailed analysis can be found in Supplementary Notes S6 and Fig S17c). In summary, the above results indicated that MONN can correctly capture the preferred interaction partners for different types of atoms in the compounds, according to the possibility of forming different kinds of non-covalent interactions.

## 4 Conclusion

Accurately predicting compound-protein interactions can greatly facilitate the drug discovery process. While several deep learning based tools have been proposed to predict binding affinities and improve virtual high-throughput screening, our approach MONN goes further to explore more about the mechanisms underlying CPIs. In this work, we demonstrated that MONN can successfully predict the pairwise non-covalent interaction matrices, which can also be used to infer the interaction sites in compounds and proteins. Comparison tests showed that MONN can outperform other state-of-the-art machine learning methods in predicting binding affinities. Besides, the structure-free input of MONN allows it to have a wider range of applications than those structure-dependent approaches. We also verified that the predictions of MONN are accordant with chemical rules, in terms of the correlation in hydrophobicity between interaction sites in compounds and proteins, and the preference of interaction partners for different atom types. All these results indicated that MONN can provide a powerful and useful tool to advance the drug development process.

## Acknowledgements

The authors thank Dr. Hailin Hu for helpful discussions about this work, and Mr. Tingzhong Tian for helpful suggestions about the manuscript.

