## Supplementary Notes and Figures for "MONN: a Multi-Objective Neural Network for Predicting Pairwise Non-Covalent Interactions and Binding Affinities between Compounds and Proteins"

### S1 Model implementation and training

#### S1.1 The graph warp unit of the graph convolution module

Both message passing unit [1] and graph warp unit [2] are involved in each iteration of the graph convolution module of MONN. We first explain the details about the graph warp unit, which executes information sharing between the atoms and the additional super node. The a super node is a virtual node introduced to encode the global feature of a compound. Before all the graph convolution iterations, the super node feature $\boldsymbol{s}^0 \in \mathbb{R}^{h_1}$ is initialized as the summation of all the atom features, that is,

$$\boldsymbol{s}^0 = \sum_{i=1}^{N_a} \boldsymbol{v}_i^0. \quad (1)$$

In the $l$-th iteration, the message passing unit first updates the atom features to obtain $\{\boldsymbol{u}_i^l\}_{i=1}^{N_a}$, as described in the Methods section of the main text. Accordingly, the graph warp unit first updates the super node feature by a single-layer neural network to obtain $\boldsymbol{u}_s^l$, that is,

$$\boldsymbol{u}_s^l = \tanh(\boldsymbol{W}_{super}^l \boldsymbol{s}^{l-1}), \tag{2}$$

where $l = 1, 2, ..., L$, $\tanh(\cdot)$ stands for the hyperbolic tangent activation function, $\boldsymbol{s}^{l-1}$ stands for the super node feature from the $(l-1)$-th iteration, and $\boldsymbol{W}_{super}^l \in \mathbb{R}^{h_1 \times h_1}$ denotes learnable parameters. Note that for all the single-layer neural networks described in Supplementary Notes, the bias terms are omitted for clarity, and $\boldsymbol{W}_x$ ($x$ can be any subscripts) denotes the learnable weight parameter.

Then, three steps are conducted to obtain the updated atom and super node features for this iteration.

Step 1: gathering information from the super node and the main nodes (atoms). The information ($\boldsymbol{u}_{s \to v}^l$) gathered from the super node is calculated by a single-layer neural network, that is,

$$\boldsymbol{u}_{s \to v}^l = \tanh(\boldsymbol{W}_{s \to v}^l \boldsymbol{s}^{l-1}), \tag{3}$$

where $l = 1, 2, ..., L$ and $\boldsymbol{W}_{s \to v}^l \in \mathbb{R}^{h_1 \times h_1}$.

To calculate the information gathered from each atom (main node), attention mechanism is used to weigh the contributions of individual atoms. As in the original version of graph warp unit [2], a K-head attention mechanism is used to determine the contribution of features from each atom when updating the super node feature, that is,

$$\boldsymbol{u}_{v \to s}^l = \tanh(\boldsymbol{W}_{v \to s}^l [\sum_{i=1}^{N_a} \alpha_{v,i}^{1,l} \boldsymbol{v}_i^{l-1}, \sum_{i=1}^{N_a} \alpha_{v,i}^{2,l} \boldsymbol{v}_i^{l-1}, ..., \sum_{i=1}^{N_a} \alpha_{v,i}^{K,l} \boldsymbol{v}_i^{l-1}]), \tag{4}$$

$$\alpha_{v,i}^{k,l} = \text{softmax}(\boldsymbol{W}_{att}^{k,l} \boldsymbol{b}_{v,i}^{k,l}), \quad k = 1, 2, ..., K, i = 1, 2, ..., N_a, \tag{5}$$

$$\boldsymbol{b}_{v,i}^{k,l} = \tanh(\boldsymbol{W}_{vatt}^{k,l} \boldsymbol{v}_i^{l-1}) * \tanh(\boldsymbol{W}_{satt}^{k,l} \boldsymbol{s}^{l-1}), \quad k = 1, 2, ..., K, i = 1, 2, ..., N_a, \tag{6}$$

where $l = 1, 2, ..., L,$ , $\boldsymbol{W}^l_{v\to s} \in \mathbb{R}^{h_1\times Kh_1}, \boldsymbol{W}^{k,l}_{att} \in \mathbb{R}^{1\times h_1}, \boldsymbol{W}^{k,l}_{vatt}$ and $\boldsymbol{W}^{k,l}_{satt} \in \mathbb{R}^{h_1\times h_1}$, $\mathrm{softmax}(x_i) = \frac{\exp(x_i)}{\sum_i \exp(x_i)}$ stands for the softmax normalization function, $[\cdot, \cdot, ..., \cdot]$ denotes the concatenation operation, $*$ denotes the element-wise multiplication, and $K$ is the number of heads.

Step 2: calculating the passed information using warp gates. For the super node, an element-wise warp gate $\boldsymbol{g}^l_{v\to s} \in \mathbb{R}^{h_1}$ is used to combine the information from the super node itself $\boldsymbol{u}^l_s$ and the main nodes (atoms) $\boldsymbol{u}^l_{v\to s}$, that is,

$$\boldsymbol{g}^l_{v\to s} = \sigma(\boldsymbol{W}^l_{gate11}\boldsymbol{u}^l_{v\to s} + \boldsymbol{W}^l_{gate12}\boldsymbol{u}^l_s), \tag{7}$$

$$\boldsymbol{t}^l_{v\to s} = (\boldsymbol{1} - \boldsymbol{g}^l_{v\to s}) * \boldsymbol{u}^l_{v\to s} + \boldsymbol{g}^l_{v\to s} * \boldsymbol{u}^l_s, \tag{8}$$

where $l = 1, 2, ..., L,$ $\boldsymbol{W}^l_{gate11}, \boldsymbol{W}^l_{gate12} \in \mathbb{R}^{h_1\times h_1}$, $\sigma(\cdot)$ stands for the sigmoid activation function, $\boldsymbol{1}$ is an all-one vector of length $h_1$, and $\boldsymbol{t}^l_{v\to s}$ denotes the information passed to super node.

For each atom, similarly, an element-wise warp gate $\boldsymbol{g}^l_{s\to i} \in \mathbb{R}^{h_1}$ is used to combine the updated atom features $\boldsymbol{u}^l_i$ and the information from the super node $\boldsymbol{u}^l_{s\to v}$, that is,

$$\boldsymbol{g}^l_{s\to i} = \sigma(\boldsymbol{W}^l_{gate21}\boldsymbol{u}^l_i + \boldsymbol{W}^l_{gate22}\boldsymbol{u}^l_{s\to v}), \tag{9}$$

$$\boldsymbol{t}^l_{s\to i} = (\boldsymbol{1} - \boldsymbol{g}^l_{s\to i}) * \boldsymbol{u}^l_i + \boldsymbol{g}^l_{s\to i} * \boldsymbol{u}^l_{s\to v}, \tag{10}$$

where $l = 1, 2, ..., L,$ $i = 1, 2, ..., N_a,$ $\boldsymbol{W}^l_{gate21}, \boldsymbol{W}^l_{gate22} \in \mathbb{R}^{h_1\times h_1}$, and $\boldsymbol{t}^l_{s\to i}$ denotes the information passed to each atom.

Step 3: calculating the updated features using gated recurrent units (GRUs) [3]. Here, two GRUs are used to determine the proportion of information updated at layer $l$ for the atom and super node features, that is,

$$\boldsymbol{v}^l_i = \mathrm{GRU}_{\mathrm{v}}(\boldsymbol{v}^{l-1}_i, \boldsymbol{t}^l_{s\to i}), i = 1, 2, ..., N_a, \tag{11}$$

$$\boldsymbol{s}^l = \mathrm{GRU}_{\mathrm{s}}(\boldsymbol{s}^{l-1}, \boldsymbol{t}^l_{v\to s}). \tag{12}$$

After completing in total $L$ iterations of graph convolution, we finally obtain a set of updated atom features $\{\boldsymbol{v}_i^L \in \mathbb{R}^{h_1}\}_{i=1}^{N_a}$ and a super node feature $\boldsymbol{s}^L \in \mathbb{R}^{h_1}$ for the input compound.

#### S1.2 Implementation of the affinity prediction module

As described in the Methods section of the main text, compound and protein attentions are employed in the affinity prediction module to reduce the atom and residue features into fixed-size vector representations. Here, we modified the dual attention network (DAN) [4] to iteratively generate the compound and protein attentions.

Before all the DAN iterations, we first define the initial compound feature $\boldsymbol{h}_c^0 \in \mathbb{R}^{h_2}$, the initial protein feature $\boldsymbol{h}_p^0 \in \mathbb{R}^{h_2}$ and the initial memory vector $\boldsymbol{m}^0 \in \mathbb{R}^{h_2}$ ($h_2$ is the hidden size used in the affinity prediction module), that is,

$$\boldsymbol{h}_c^0 = \frac{1}{N_a}\sum_{i=0}^{N_a} \boldsymbol{h}_{v,i}, \tag{13}$$

$$\boldsymbol{h}_p^0 = \frac{1}{N_r}\sum_{j=0}^{N_r} \boldsymbol{h}_{r,j}, \tag{14}$$

$$\boldsymbol{m}^0 = \boldsymbol{h}_c^0 * \boldsymbol{h}_p^0, \tag{15}$$

where $\{\boldsymbol{h}_{v,i}\}_{i=1}^{N_a}$ and $\{\boldsymbol{h}_{r,j}\}_{j=1}^{N_r}$ are the transformed atom and residue features, as also described in the main text.

The compound feature, protein feature and the memory vector are then updated by $D$ iterations of DAN. Specifically, at the $d$-th iteration ($d = 1, 2, ..., D$), we first calculate the information shared between the atom features and the residue features, according to the predicted pairwise interaction matrix $\boldsymbol{P}$. Intuitively, each atom of the compound receives information from all the residues of the protein, weighed by the probability of local interactions between the atom-residue pairs, and vice versa for the residues. That is,

$$\boldsymbol{s}^d_{r\to v,i} = \sum_{j=1}^{N_r} P_{ij}\tanh(\boldsymbol{W}^d_{r\to v}\boldsymbol{h}_{r,j}), i = 1, 2, ..., N_a, \tag{16}$$

$$\boldsymbol{s}^d_{v\to r,j} = \sum_{i=1}^{N_a} P_{ij}\tanh(\boldsymbol{W}^d_{v\to r}\boldsymbol{h}_{v,i}), j = 1, 2, ..., N_r, \tag{17}$$

where $d = 1, 2, ..., D$, $\boldsymbol{W}^d_{r\to v}, \boldsymbol{W}^d_{v\to r} \in \mathbb{R}^{h_2\times h_2}$, $\{\boldsymbol{s}^d_{r\to v,i}\}_{i=1}^{N_a}$ are the information delivered from residues to the atoms, and $\{\boldsymbol{s}^d_{v\to r,j}\}_{j=1}^{N_r}$ are the information delivered from atoms to the residues.

Next, the atom or residue features ($\{\boldsymbol{h}^d_{v,i}\}_{i=1}^{N_a}$ or $\{\boldsymbol{h}^d_{r,j}\}_{j=1}^{N_r}$), the memory vector from the previous iteration ($\boldsymbol{m}^{d-1}$) and the above derived shared information ($\{\boldsymbol{s}^d_{r\to v,i}\}_{i=1}^{N_a}$ or $\{\boldsymbol{s}^d_{v\to r,j}\}_{j=1}^{N_r}$) are combined to calculate the hidden states of the compound and protein attentions, that is,

$$\boldsymbol{b}^d_{v,i} = \tanh(\boldsymbol{W}^d_{vc}\boldsymbol{h}_{v,i}) * \tanh(\boldsymbol{W}^d_{mc}\boldsymbol{m}^{d-1}) * \boldsymbol{s}^d_{r\to v,i}, \tag{18}$$

$$\boldsymbol{b}^d_{r,j} = \tanh(\boldsymbol{W}^d_{rp}\boldsymbol{h}_{r,j}) * \tanh(\boldsymbol{W}^d_{mp}\boldsymbol{m}^{d-1}) * \boldsymbol{s}^d_{v\to j}, \tag{19}$$

where $d = 1, 2, ..., D$ and $\boldsymbol{W}^d_{vc}, \boldsymbol{W}^d_{mc}, \boldsymbol{W}^d_{rp}, \boldsymbol{W}^d_{mp} \in \mathbb{R}^{h_2\times h_2}$.

The compound and protein attentions are then calculated through two linear layers, normalized by the softmax function, that is,

$$\alpha^d_{v,i} = \text{softmax}(\boldsymbol{W}^d_{vs}\boldsymbol{b}^d_{v,i}), \tag{20}$$

$$\alpha^d_{r,j} = \text{softmax}(\boldsymbol{W}^d_{rs}\boldsymbol{b}^d_{r,j}), \tag{21}$$

where $d = 1, 2, ..., D$ and $\boldsymbol{W}^d_{vs}, \boldsymbol{W}^d_{rs} \in \mathbb{R}^{1\times h_2}$.

Finally, the fixed-size compound feature, protein feature and memory vector for the

$d$-th iteration is updated by current attentions:

$$\boldsymbol{h}_c^d = \sum_{i=0}^{N_a} \alpha_{v,i}^d \boldsymbol{h}_{v,i}, \tag{22}$$

$$\boldsymbol{h}_p^d = \sum_{j=0}^{N_r} \alpha_{r,j}^d \boldsymbol{h}_{r,j}, \tag{23}$$

$$\boldsymbol{m}^d = \mathrm{GRU}(\boldsymbol{m}^{d-1}, \boldsymbol{h}_c^d * \boldsymbol{h}_p^d), \tag{24}$$

where GRU is the gated recurrent unit [3].

After completing all the $D$ iterations, we obtain the final compound attention $\{\alpha_{v,i}^D\}_{i=1}^{N_a}$ and protein attention $\{\alpha_{r,j}^D\}_{j=1}^{N_r}$. In the main text, we drop the superscript $D$ for simplicity and use $\{\alpha_{v,i}\}_{i=1}^{N_a}$ and $\{\alpha_{r,j}\}_{j=1}^{N_r}$ to denote the final compound and protein attentions, respectively.

### S2 Training and hyper-parameter calibration

#### S2.1 Clustering-based cross validation

##### S2.1.1 Clustering

In the real datasets for compound-protein interaction prediction, there often exist highly similar compounds or proteins. To avoid the data redundancy problem caused by these similar compounds or proteins, we follow the same strategy as in [5] and use a clustering-based cross validation strategy to evaluate the performance of our prediction model. The train-test splitting process in such a clustering-based cross validation scheme guarantees that the compounds (or proteins) within the same cluster, which share high similarities, are either all used in the training set, or all used in the test set. Note that an alternative approach to reduce data redundancy is to discard those high-similar data points. However, we argue that the clustering-based scheme would allow us to make better use of all the available data. We use the single-linkage clustering algorithm [6], which ensures that the minimal distance between any two clusters is above a given distance threshold. Here, the distance between a pair of compounds $(c_i, c_j)$, is defined as

$$\text{Distance}(c_i, c_j) = 1 - \text{Jaccard}(\text{MF}(c_i), \text{MF}(c_j)), \tag{25}$$

where MF(·) is the Morgan fingerprints calculated by RDKit [7] and Jaccard(·, ·) denotes the Jaccard similarity.

The distance between a pair of proteins $(p_i, p_j)$ is defined as

$$\text{Distance}(p_i, p_j) = 1 - \frac{SW(p_i, p_j)}{\sqrt{(\text{SW}(p_i, p_i)\text{SW}(p_j, p_j))}}, \tag{26}$$

where SW(·, ·) stands for the Smith-Waterman alignment score calculated based on the SSW library (https://github.com/mengyao/Complete-Striped-Smith-Waterman-Library). The distance threshold values used in our paper is [0.3, 0.4, 0.5, 0.6]. We choose 0.3 as the lower limit of the threshold, because a distance smaller than 0.3 would not be separable enough to avoid the data redundancy problem, consistent with the previous study [5]. The upper limit of our clustering threshold is set to 0.6, because a higher threshold will lead to so large clusters that the splitting of training-test data would be highly imbalanced (*i.e.*, too much training data and too little test data, or vice versa, Tables S1,S2).

Table S1: Sizes of compound clusters under different thresholds. The total number of compounds is 10257.

| **Threshold** | **Cluster number** | **Max cluster size** |
|---|---|---|
| 0.1 | 9894 | 9 |
| 0.2 | 9317 | 48 |
| 0.3 | 7990 | 98 |
| 0.4 | 6467 | 318 |
| 0.5 | 4581 | 1421 |
| 0.6 | 2473 | 5659 |
| 0.7 | 438 | 9659 |
| 0.8 | 10 | 10246 |
| 0.9 | 1 | 10257 |

Table S2: Sizes of protein clusters under different thresholds. The total number of proteins is 2577.

| **Threshold** | **Cluster number** | **Max cluster size** |
|---|---|---|
| 0.1 | 2325 | 22 |
| 0.2 | 2248 | 24 |
| 0.3 | 2163 | 24 |
| 0.4 | 2060 | 47 |
| 0.5 | 1918 | 53 |
| 0.6 | 1716 | 55 |
| 0.7 | 1405 | 56 |
| 0.8 | 69 | 2462 |
| 0.9 | 1 | 2577 |

##### S2.1.2 Cross validation settings

After generating the compound and protein clusters, three settings are considered during the cross validation process, *i.e.*, the new-compound setting, the new-protein setting and the both-new setting. To explain these settings, we denote the training, validation and test sets by $D_{train}, D_{valid}$ and $D_{test}$, respectively, and use $(c_i, p_i)$ to represent the compound-protein pair of the $i$-th sample ($i = 1, 2, ..., N$).

In the new-compound setting, cross validation is performed on compound clusters, so that the compound-protein pairs with compounds from the same cluster cannot be shared across training, valid and test sets. That is, for any two compound-protein pairs $(c_i, p_i)$ and $(c_j, p_j)$ from different sets, $c_i$ and $c_j$ must come from different compound clusters.

In the new-protein setting, cross validation is performed on protein clusters, so that the compound-protein pairs with proteins from the same cluster cannot be shared across training, valid and test sets. That is, for any two compound-protein pairs $(c_i, p_i)$ and $(c_j, p_j)$ from different sets, , $p_i$ and $p_j$ must come from different protein clusters.

In the both-new setting, both compound clusters and protein clusters cannot be shared across training, valid and test sets. That is, for any two compound-protein pairs $(c_i, p_i)$ and $(c_j, p_j)$ from different sets, , $c_i$ and $c_j$ must come from different compound clusters, and $p_i$ and $p_j$ must come from different protein clusters.

For the new-compound and the new-protein settings, we use five-fold cross validation, and the train-valid-test splitting ratio is approximately 7 : 1 : 2. Note that here the ratio is an approximation, because the splitting is performed on clusters, and the number of data points among individual clusters is not necessarily evenly distributed. For the both-new setting, we randomly partition the pairs of compound-protein clusters into a $3 \times 3$ grid (Fig S8). Then, a nine-fold cross validation [8] was conducted according to the following three steps: 1) select a grid as the test set; 2) discard the four grids that share compound or protein clusters with the selected one; 3) reorganize the remaining grids as a new $3 \times 3$ grid setting and randomly select one grid as the validation set, and the four grids that do not share any compound or protein clusters with the validation set are used as the training set. Such a cross-validation strategy results in an approximately 16 : 4 : 9 train-valid-test ratio.

#### S2.2 Hyper-parameter selection

Four baseline models were included in the performance comparison for the binding affinity prediction task: CGKronRLS [9], DeepDTA [10], Tsubaki el al.'s method [11] and DeepAffinity [12]. Gao et al.'s method [13] was not included here, because the source code was not released, and the model requires additional input information (*i.e.*, GO terms of proteins). For our model and all the baseline methods, each cross-validation setting (*i.e.*, new-compound, new-protein or both-new) has a specific set of hyper-parameters. For MONN, the hyper-parameter selection was performed with both training objectives. The details of the hyper-parameter spaces for MONN and the baseline methods are listed below:

- For our model, the number of graph convolution layers $L = 4$ and the number of DAN iterations $D = 2$ were determined using the same schemes as in the original papers [1,4]. The hidden size $h_1$ is set to 128. Other hyper-parameters include the number of heads of the k-head attention used in the graph convolution module $K \in \{1, 2\}$, the number of CNN layers $L_{CNN} \in \{2, 4\}$, the kernel size of the CNN layers $S_{kernel} \in \{5, 7\}$, the hidden size of the affinity prediction module $h_2 \in \{64, 128\}$, the ratio of pairwise loss $\lambda \in \{0, 0.1, 1\}$. A grid search was used to search for the best combination of these four hyper-parameters.

- For CGKronRLS [8], the regularization parameter was chosen from $\{2^{-5}, 2^{-4}, ..., 2^{5}\}$.

- For DeepDTA [10], a grid search was conducted to select the best combination of different hyper-parameters, including the number of filters from $\{16, 32, 64, 128\}$, the length of sequence windows from $\{4, 8, 12\}$ and the length of smiles windows from $\{4, 6, 8\}$. These ranges were adapted from the original paper [10].

- For Tsubaki et al.'s method [11], according to the original paper, the radius of compound subgraph and the length of the protein "ngram" (*i.e.*, n-mer from protein sequence) were selected from $\{(0, 1), (1, 2), (2, 3)\}$, the hidden dimension of ngram and atom embedding was selected from $\{5, 10, 20, 30\}$, and the number of layers of both CNN and GNN was selected from $\{2, 3, 4\}$. A grid search was performed to select the best combination from these ranges. Note that Tsubaki et al.'s method is originally a classification model. Here, we modified its last hidden layer by removing the activation function, and changed its loss function to the mean squared error (MSE) to perform the regression task.

- For DeepAffinity [12], since the authors did not provide specific hyper-parameter set and their model requires a pretrain step, we directly used their pretrained RNN-CNN models and then fine-tuned them with our data. For each cross-validation setting, we chose the better DeepAffinity model from models with either joint or separate attentions.

Apart from all the hyper-parameters mentioned above, all the methods have another hyper-parameter, *i.e.*, the number of epochs (or iterations) for the training process. We used the RMSE as the evaluation metric from the validation set to select the best value of this hyper-parameter for all the methods. The maximum number of epochs for our method was set to 30. For DeepDTA, Tsubaki et al.'s method and DeepAffinity, we used their default maximum numbers of epochs (which is 100). For CGKronRLS, we set the maximum number of iterations to 500, as the performance no longer increased after 500 iterations.

For each cross-validation setting, the best hyper-parameters were selected by the IC50 dataset with clustering threshold 0.3. The same parameters were used for other scenarios (*i.e.*, other thresholds and the KIKD dataset) under the corresponding cross-validation setting. For pairwise interaction prediction, we also used the best hyper-parameters selected based on the affinity prediction results. We did not select the hyper-parameters according to the performance of pairwise prediction task (that is, only single training objective of MONN was used), for the following two reasons: 1) For MONN, the affinity prediction task involves all the modules of our model; 2) For baseline methods, they do not include a direct supervised optimization procedure for local interaction prediction, so for fair comparison, we did not specifically tune the hyper-parameters for pairwise interaction prediction for either MONN or the baseline methods.

### S3 Supplementary details on dataset construction

#### S3.1 Supplementary details on the construction of the benchmark dataset

We downloaded the protein-ligand complexes and their corresponding binding affinity data from the PDBbind database (version 2018, the general set, http://www.pdbbind-cn.org/index.asp). Each complex was provided with an affinity value of certain measurement type (*e.g.*, $\mathrm{K_i}$, $\mathrm{K_d}$, or $\mathrm{IC_{50}}$). To prepare the training labels of the pairwise non-covalent interactions, we also downloaded the structure data files (.pdb) from the RCSB PDB (https://www.rcsb.org/) according to the PDB IDs provided in the PDBbind. These structure files were then processed by PLIP [14] to extract the non-covalent interactions and generate the pairwise interaction labels. The detailed process of dataset construction is described below.

There are in total 16,151 entries in the downloaded PDBbind dataset [15, 16]. We filtered these entries according to the following criteria: 1) affinity values need to be an accurate number, rather than a range or an approximation; 2) compounds need to have available and valid graph representations that can be processed by RDKit [7]; and 3) proteins need to be successfully mapped to UniProt IDs with available sequence data. Note that we used UniProt sequences instead of the protein sequences directly extracted from the PDB structures, for the following reasons: first, the protein sequences in the PDB structures may be incomplete (*e.g.*, only include some domains or lack some flexible regions); second, one protein may have different sequence variants in different PDB structures; third, in a practical scenario, when we want to predict candidate ligands for a protein without known structure, it is more convenient to use its full-length primary sequence. In total, we obtained 13,306 compound-protein pairs satisfying those criteria.

Next, we calculated the pairwise interaction labels for the resulting 13,306 compound-protein pairs. The non-covalent interactions between the proteins and the corresponding ligands were extracted by the PLIP tool [14] (https://github.com/ssalentin/plip/). The ligand atoms involved in the non-covalent interactions were then mapped to the corresponding compound structures (downloaded from http://ligand-expo.rcsb.org/ld-download.html), which contained the unique names and indices for all the non-hydrogen atoms. For proteins, the residues involved in non-covalent interactions were first mapped to the UniProt sequences using a sequence alignment tool (https://github.com/mengyao/Complete-Striped-Smith-Waterman-Library/). Then, we examined the mappings to control the quality of the generated interaction labels, and discarded those structures when the detected interactions cannot be correctly mapped into the molecular graphs and the protein sequences. In addition, to further improve the mapping quality, we also filtered the complexes whose protein sequences in the PDB structures and the corresponding UniProt sequences had less than 90% matched residues. After the mapping process, we filled the pairwise interaction matrix according to the indices of the atoms and residues involved in the non-covalent interactions to obtain the final interaction labels. After these procedures, we successfully constructed pairwise non-covalent interaction labels for about 95% of the compound protein pairs, resulting in 12,738 interaction matrices out of the 13,306 complex structures.

After constructing the dataset as described above, the performance of the pairwise interaction prediction was evaluated using all the available data. For binding affinity prediction, we further separated the compound-protein pairs according to the measurement types of binding affinity (*i.e.*, $\mathrm{K_i}, \mathrm{K_d}$ or $\mathrm{IC_{50}}$), resulting in two affinity datasets, which were called the IC50 dataset and the KIKD dataset, respectively. The reason for this separation was that the $\mathrm{IC_{50}}$ values usually depend on the experimental conditions and thus are often considered to be noisier. Therefore, here we mainly used the IC50 dataset for hyper-parameter tuning for binding affinity prediction. For those repetitive records (defined as pairs with the same protein ID and the same compound InChI), we only kept the pairs with pairwise interaction labels and with higher binding affinity. Finally, we obtained 5,340 and 6,689 unique pairs for the IC50 and KIKD datasets, respectively.

#### S3.2 Construction of the additional test dataset for validating pairwise non-covalent interaction predictions

We also downloaded the compound-protein complexes from the RCSB PDB database [17] to generate an additional test set for evaluating the pairwise non-covalent interaction prediction results of MONN. Since the PDBbind v2018 dataset, which was used as our training data, already contained the high-quality compound-protein complex structures with releasing date up to the end of 2017, we downloaded structure data with date from January, 2018 to March, 2019 to avoid overlap between training and additional test datasets. Here three criteria were used to select the compound-protein complexes and control the quality of this additional dataset: (1) the protein sequences can be mapped to a Uniprot sequence, with at least 90% perfect matches in sequence alignment; (2) to remove ions, coenzymes and other crystallization assistant chemicals, we retained only those test compound-protein pairs in which the quantitative estimation of drug-likeness (QED) scores [18] of the compounds are larger than 0.5; and (3) overlaps between training and test datasets were removed by discarding the test samples with both compound and protein similarities larger than 0.9 with any compound-protein pair in the training data. Then, the selected compound-protein complexes were processed by PLIP [14] (https://github.com/ssalentin/plip/) to extract the non-covalent interactions and construct the pairwise interaction labels as described in Section S3.1.

### S4 Evaluating different types of neural attentions

#### S4.1 Evaluation metrics

We used the average AUC scores and the average enrichment scores to evaluate the interpretability of neural attentions and prediction performance of MONN. Given a test dataset containing $N$ samples, the average AUC score is defined as:

$$\text{average AUC score} = \frac{1}{N}\sum_{n=1}^{N}\text{AUC}(n), \tag{27}$$

where AUC($n$) stands for the area under the ROC curve calculated between the labels and the predictions of the $n$-th sample.

The average enrichment score is defined as:

$$\text{average enrichment score} = \frac{1}{N}\sum_{n=1}^{N}\text{enrichment}(n), \tag{28}$$

$$\text{enrichment}(n) = \frac{\text{precision}(n)}{\text{random\_precision}(n)}, \tag{29}$$

where precision($n$) stands for the precision score between the true labels and binarized predictions (defined below) of the $n$-th sample, and random_precision($n$) is the expected precision of random predictions. Suppose that the positive-negative ratio of the whole dataset is $x_{pos} : x_{neg}$, and the length of prediction is $l_{pred}$. Then the binarization is realized by sorting the real-value predictions, and assigning 1 for top $\lceil l_{pred} \times x_{pos}/(x_{pos}+x_{neg}) \rceil$ predictions ($\lceil \cdot \rceil$ is the ceiling operation), and 0 for the rest. The random_precision($n$) is calculated as random_precision($n$) $= x_{pos}/(x_{pos}+x_{neg})$.

The upper limit of the average enrichment score is derived below:

$$\begin{aligned} \text{average enrichment score} &= \frac{1}{N}\sum_{n=1}^{N}\text{enrichment}(n) \\ &= \frac{1}{N}\sum_{n=1}^{N}\frac{\text{precision}(n)}{x_{pos}/(x_{pos}+x_{neg})} \\ &\leq \frac{1}{N}\sum_{n=1}^{N}\frac{1}{x_{pos}/(x_{pos}+x_{neg})} \\ &= 1+\frac{x_{neg}}{x_{pos}}. \end{aligned} \tag{30}$$

Thus with a relatively small positive-negative ratio (*i.e.*, relatively large $\frac{x_{neg}}{x_{pos}}$), the upper limit of the average enrichment score is relatively high.

#### S4.2 Implementation of the tested neural attentions

We tested four types of neural attentions, by either using the original implementation or re-implementing and incorporating them into our MONN framework. For the protein attentions in the Tsubaki et al.'s method [11], we directly used their source code. Since only the attention for proteins is generated, evaluations in terms of compound interaction sites and pairwise interactions are not applicable for the Tsubaki et al.'s method. For the bilinear attention in Gao et al.'s method [13], since the source code was not released, we implemented it according to the description in the original paper. For the separate attention and joint attention of DeepAffinity [12], their original implementation of attentions was used to weigh short secondary protein structures (SPSs), rather than single residues. Thus we re-implemented their attentions for testing them in our condition, which requires the protein attentions to be calculated at residue resolution.

In our implementations, we used the transformed atom and residue features ($\{\boldsymbol{h}_{v,i}\}_{i=1}^{N_a}$

and $\{\boldsymbol{h}_{r,j}\}_{j=1}^{N_r}$) from the affinity prediction module of MONN (described in the main text) to calculate the compound and protein attentions according to the neural attention based methods mentioned above. After that, the resulting compound and protein attentions substitute the corresponding part (*i.e.*, the DAN part) in our affinity prediction module, and then the models were trained according to the binding affinity labels. Note that our pairwise interaction prediction module is not used in this process. More details about how we implemented these neural attentions under our MONN's framework are described below.

##### S4.2.1 The bilinear attention of Gao et al.'s method

First, the atom features and the residue features are combined to calculate a soft alignment matrix $\boldsymbol{P}$ of size $N_a \times N_r$:

$$P_{ij} = \tanh((\boldsymbol{W}_{uv}\boldsymbol{h}_{v,i})^T(\boldsymbol{W}_{ur}\boldsymbol{h}_{r,j})), \tag{31}$$

where $\boldsymbol{W}_{uv}, \boldsymbol{W}_{ur} \in \mathbb{R}^{h_2 \times h_2}$ are learnable weight parameters.

Then, the compound attentions $\{\alpha_{v,i}\}_{i=1}^{N_a}$ and protein attentions $\{\alpha_{r,j}\}_{j=1}^{N_r}$ are calculated by max-pooling over the soft alignment matrix $P$, followed by a softmax normalization function:

$$\alpha_{v,i} = \text{softmax}(\max_{j=1,2,\ldots,N_r} P_{ij}), \tag{32}$$

$$\alpha_{r,j} = \text{softmax}(\max_{i=1,2,\ldots,N_a} P_{ij}), \tag{33}$$

where $\text{softmax}(x_i) = \frac{\exp(x_i)}{\sum_i \exp(x_i)}$ stands for the normalization function.

These attentions are then used for reducing the compound and protein features for predicting binding affinity values, as described in the Methods section of the main text. To evaluate the interpretability, the compound attentions $\{\alpha_{v,i}\}_{i=1}^{N_a}$ and protein attentions $\{\alpha_{r,j}\}_{j=1}^{N_r}$ are used as the predictions of interaction sites of compounds and proteins, respectively. The soft alignment matrix $\boldsymbol{P}$ is used as the predicted pairwise interaction matrix.

##### S4.2.2 The separate attention of DeepAffinity

The separate attention of DeepAffinity [12] calculates the soft self-attention for atoms and proteins, separately. In particular, the attentions for atoms ($\{\alpha_{v,i}\}_{i=1}^{N_a}$) are calculated by:

$$\boldsymbol{e}_{v,i} = \tanh(\boldsymbol{W}_{ev}\boldsymbol{h}_{v,i}), \tag{34}$$

$$\alpha_{v,i} = \mathrm{softmax}(\boldsymbol{W}_{av}\boldsymbol{e}_{v,i}), \tag{35}$$

where $\boldsymbol{W}_{ev} \in \mathbb{R}^{h_2 \times h_2}$ and $\boldsymbol{W}_{av} \in \mathbb{R}^{1 \times h_2}$ are learnable weight parameters, and tanh is the hyperbolic tangent activation function.

Similarly, the attentions for residues ($\{\alpha_{r,j}\}$) are calculated by:

$$\boldsymbol{e}_{r,j} = \tanh(\boldsymbol{W}_{er}\boldsymbol{h}_{r,j}), \tag{36}$$

$$\alpha_{r,j} = \mathrm{softmax}(\boldsymbol{W}_{ar}\boldsymbol{e}_{r,j}), \tag{37}$$

where $\boldsymbol{W}_{er} \in \mathbb{R}^{h_2 \times h_2}$ and $\boldsymbol{W}_{ar} \in \mathbb{R}^{1 \times h_2}$ are learnable weight parameters.

The compound and protein attentions are then used in the affinity prediction module of MONN (as described in the Methods section of the main text). After trained by binding affinity labels, they are used as predictions of the interaction sites. Evaluation on pairwise interaction prediction is not applicable for this kind of attention, as the relationships between atoms and residues are not explored in this condition.

##### S4.2.3 The joint attention of DeepAffinity

A pairwise interaction matrix $\boldsymbol{P}$ of size $N_a \times N_r$ is first calculated through a single layer neural network that combines both atom and residue features, that is,

$$P_{ij} = \tanh((\boldsymbol{W}_{uv}\boldsymbol{h}_{v,i})^T(\boldsymbol{W}_{ur}\boldsymbol{h}_{r,j})), \tag{38}$$

where $\boldsymbol{W}_{uv}, \boldsymbol{W}_{ur} \in \mathbb{R}^{h_2 \times h_2}$ are learnable weight parameters.

Then, a softmax function is used to normalize the pairwise interaction matrix over all the elements, to obtain a $N_a \times N_r$ attention matrix $\boldsymbol{A}$, that is,

$$A_{ij} = \frac{\exp(P_{ij})}{\sum_{i=1}^{N_a}\sum_{j=1}^{N_r}\exp(P_{i,j})}. \tag{39}$$

This normalized pairwise attention matrix $\boldsymbol{A}$ can be used in the evaluation of pairwise interaction prediction. In addition, through marginalizing $\boldsymbol{A}$, we can also derive the predictions of interaction sites in compounds and proteins, that is,

$$\alpha_{v,i} = \max_{j \in 1,2,\ldots,N_r} P_{i,j}, \tag{40}$$

$$\alpha_{r,j} = \max_{i \in 1,2,\ldots,N_a} P_{i,j}. \tag{41}$$

Since the original implementation of DeepAffinity with joint attention did not define the compound-wise/protein-wise attentions, here we modified our affinity prediction module, by replacing the outer product between compound and protein features with a combined feature, which is used in DeepAffinity:

$$\boldsymbol{b}_{i,j} = \tanh(\boldsymbol{W}_{bv}\boldsymbol{h}_{v,i} + \boldsymbol{W}_{br}\boldsymbol{h}_{r,j}), \tag{42}$$

$$\boldsymbol{h} = \sum_{i=1}^{N_a}\sum_{j=1}^{N_r} A_{ij}\boldsymbol{b}_{i,j}, \tag{43}$$

where $\boldsymbol{W}_{bv}, \boldsymbol{W}_{br} \in \mathbb{R}^{h_2 \times h_2}$ are learnable weight parameters.

The final binding affinity is then predicted by:

$$a = \boldsymbol{W}_a f([\boldsymbol{s}, f(\boldsymbol{h})]), \tag{44}$$

where $f(\cdot)$ is leaky ReLU activation function, $[\cdot, \cdot]$ stands for concatenation operation, and $\boldsymbol{s}$ is the super node feature as described in Section S1.1.

### S5 Performance of MONN in predicting binding affinities on a large-scale dataset

The performance of DeepAffinity [12] on its BindingDB-derived dataset was obtained from the original paper. In addition to a “single model”, the performance of several ensemble versions of DeepAffinity (that is, averaging predictions over several single models) was also reported in [12]. In particular, one ensemble strategy was called “parameter ensemble”, *i.e.*, averaging the predictions over the last 10 epochs. The other ensemble strategy was called “parameter+NN ensemble”, that is, averaging predictions over the last 10 epochs of three networks with different hyper-parameters (*i.e.*, the sizes of the last fully-connected layers).

Among the baseline methods, CGKronRLS [8] and Tsubaki et al.’s method [11] were not included in this test. CGKronRLS needs the input of compound and protein similarity matrices, and its the space and time usage increases dramatically with the increase of dataset size (for example, loading the float32-format similarity matrix of all the 202,169 unique compounds from the BindingDB training set needs 149 Gb memory, and processing such a matrix by the CGKronRLS algorithm is nearly infeasible). Unlike other deep learning-based baseline methods that process batches of input samples, Tsubaki et al.’s implementation allows only one sample to be processed at a time, so training this method on such a large dataset would be too time-consuming and nearly impractical.

For MONN and DeepDTA [10], we used the same training and test sets as provided by DeepAffinity [12], but dropped out a small number of samples (49 out of 263,583 training samples and 26 out of 113,168 test samples, about 0.02%), since these SMILES strings cannot be converted into a valid molecular graph by RDKit [7]. Followed the same strategy as used in DeepAffinity [12], 10% of training data were used as the validation set. As we used this validation set to select the best epoch for MONN and DeepDTA, the "parameter ensemble" strategy is not suitable for these two models. So we directly use 30 ensemble models for MONN and DeepDTA (the same number of DeepAffinity predictions in their "parameter+NN ensemble" setting). That is, the predictions of 30 models were calculated and averaged as the ensemble prediction. Here, we did not specifically optimize the hyper-parameters of MONN and DeepDTA over the BindingDB dataset, and directly use the three hyper-parameter settings derived from the hyper-parameter selection for the PDBbind-derived benchmark dataset (Supplementary Notes S2.2).

The performances of DeepAffinity, DeepDTA and MONN were evaluated in terms of RMSE and Pearson correlation, as listed in Table S3. When evaluating the single models, MONN achieved the best Pearson correlation (0.858). For all the methods, their performance can be largely increased by using ensemble based models. Among them, the ensemble version of MONN achieved the best performance (RMSE 0.658 and Pearson correlation 0.895), which demonstrated the superiority of our method.

Table S3: Performance evaluation of different prediction approaches on the BindingDB dataset. The RMSE and Pearson correlation of DeepAffinity are adopted from the original paper [12], in which "parameter ensemble" means averaging the predictions over the last 10 epochs, and "parameter+NN ensemble" means averaging predictions over the last 10 epochs of three networks with different hyper-parameters (*i.e.*, average over 30 predictions).

| **Method** | **RMSE** | **Pearson correlation** |
|---|---|---|
| DeepAffinity (single model) | 0.74 | 0.84 |
| DeepAffinity (parameter ensemble) | 0.73 | 0.84 |
| DeepAffinity (parameter+NN ensemble) | 0.71 | 0.86 |
| DeepDTA (single model) | 0.782 | 0.848 |
| DeepDTA (ensemble of 30 models) | 0.686 | 0.886 |
| MONN (single model) | 0.764 | 0.858 |
| MONN (ensemble of 30 models) | **0.658** | **0.895** |

### S6 Supplementary details about analyzing the chemical rules of non-covalent interactions captured by MONN

As defined in the main text, the conditional likelihood score $p$(residue property=$x$|atom property=$y$) = (Number of residues $\in S(x)$ that interact with the atoms of property $y$)/(Total number of residues interacting with the atoms of property $y$), where $S(x)$ represents of the set of residues with property $x$. Here, to be more specific, $S$("H-bond donor") = {H, K, N, Q, R, S, T, W, Y}, in which all the residues have at least one hydrogen bond donor in their side chains. Similarly, $S$("H-bond acceptor") = {D, E, H, N, Q, S, T, Y}, and $S$("aromatic") = {Y, W, F}. The corresponding properties of atoms from the compounds were calculated using RDKit [7].

For $\pi$-stacking interactions, there are three amino acids, *i.e.*, phenylalanine, tryptophan and tyrosine, containing aromatic rings. They generally have higher conditional likelihood scores when their interaction partners are aromatic atoms from the compounds (0.44 calculated from true labels compared to 0.09 from random control, Fig S17c). In the MONN prediction results, the three aromatic residues also have higher conditional likelihood scores (0.35 from cross validation and 0.37 from the additional test set, Fig S17c) than that from random control, which thus provided another evidence to support the reasonableness of the MONN prediction results.

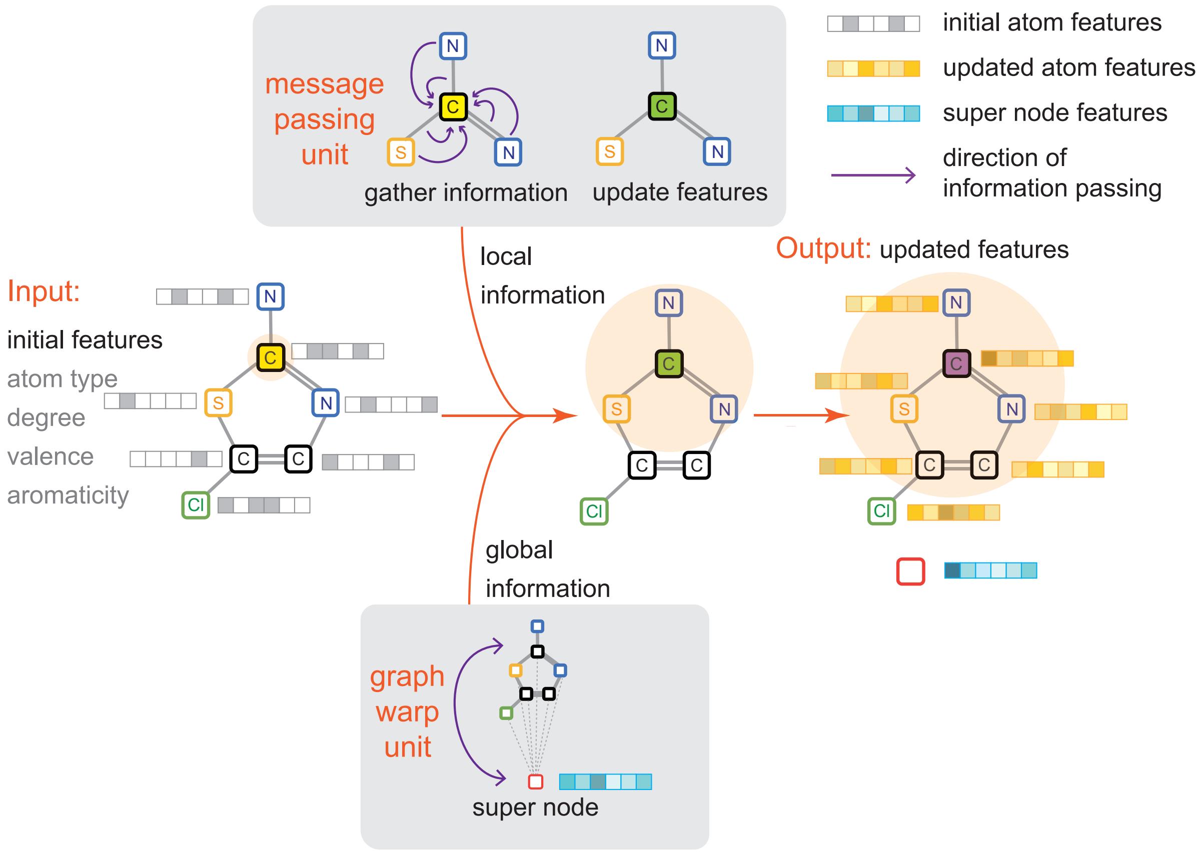


Figure S1: The graph convolution module of MONN.

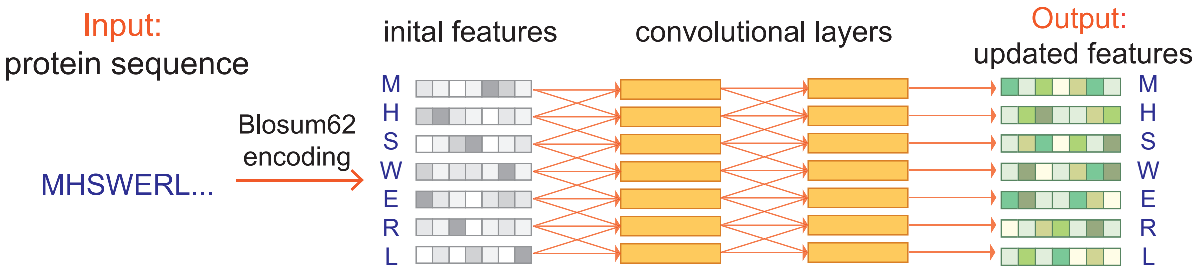


Figure S2: The convolution neural network (CNN) module of MONN.

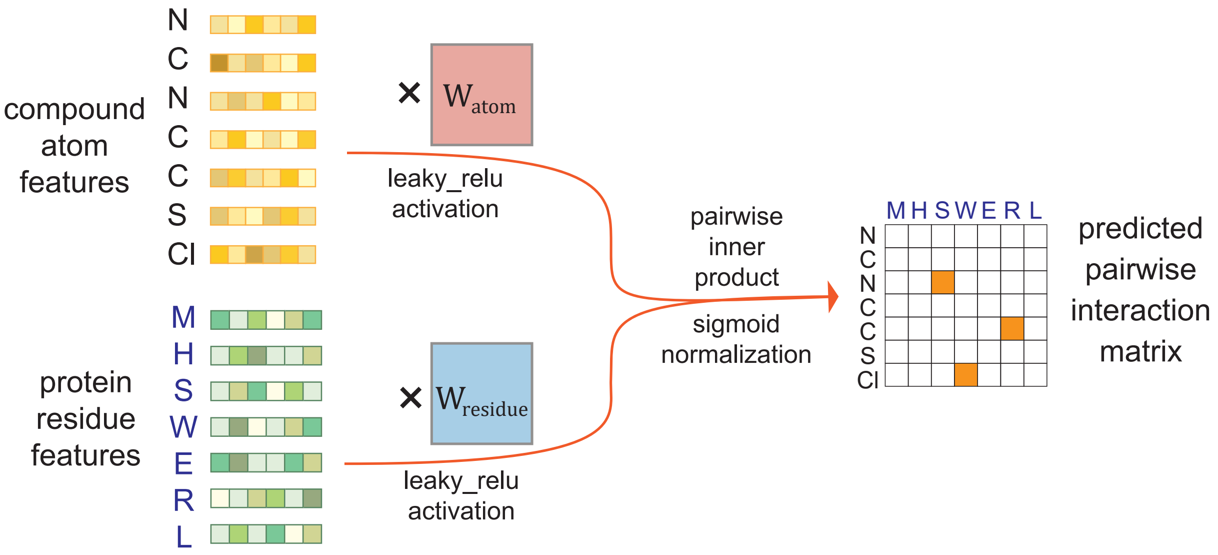


Figure S3: The pairwise interaction prediction module of MONN. Here, $W_{atom}$ and $W_{residue}$ are weight parameters of two single-layer neural networks.

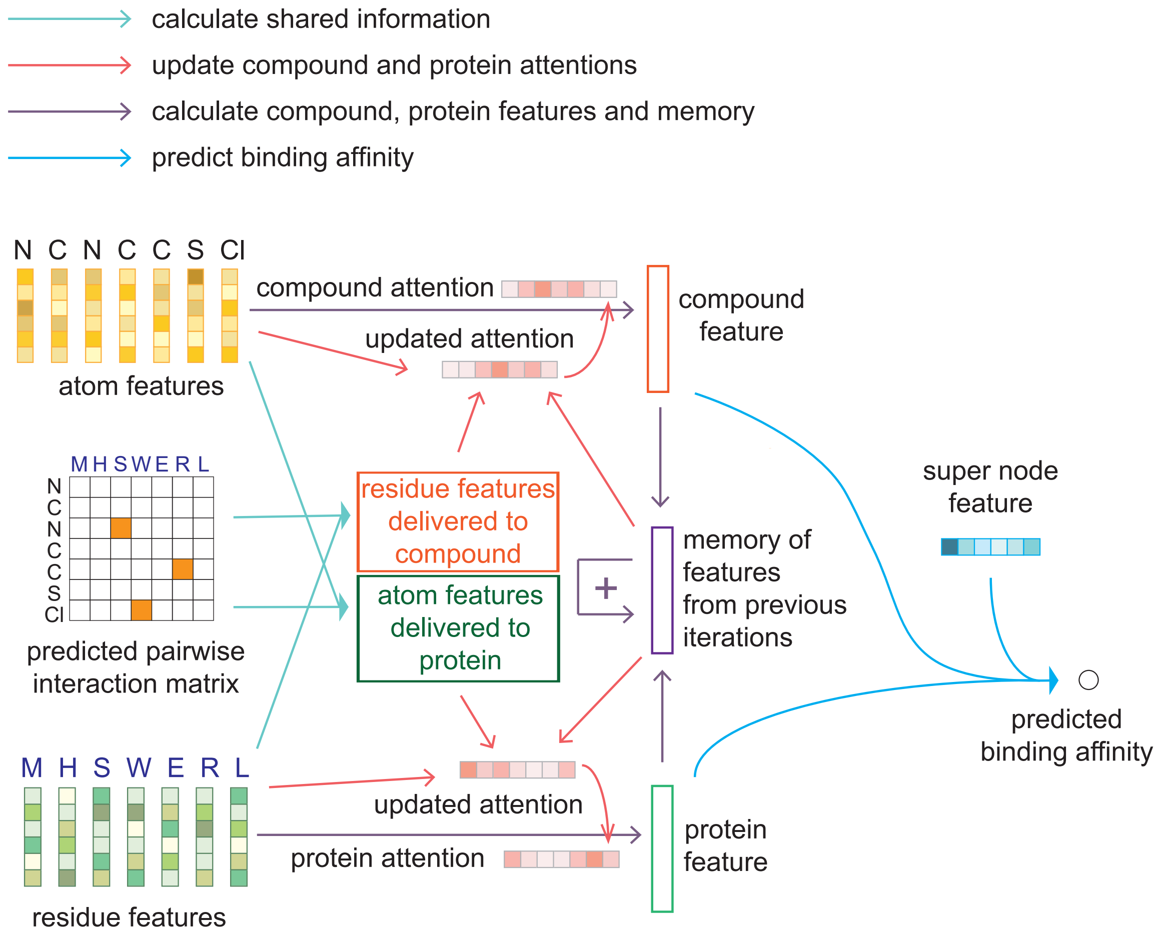


Figure S4: The affinity prediction module of MONN.

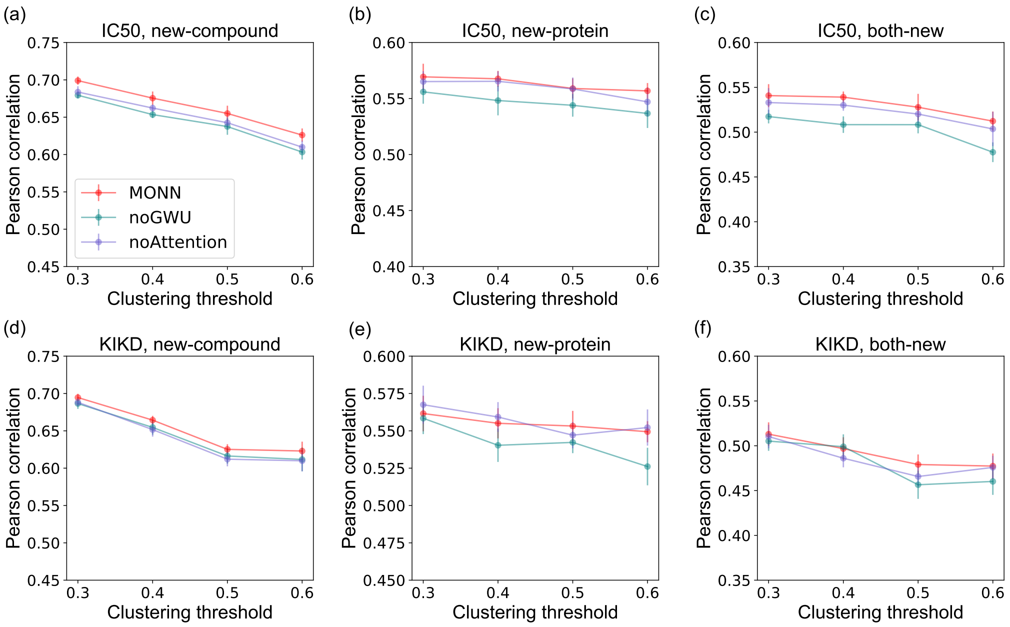


Figure S5: The ablation study for the binding affinity prediction task by removing the graph warp unit (GWU) or the attention part (DAN) of MONN. Performance was evaluated in terms of Pearson correlation, using the IC50 (a-c) and KIKD (d-f) datasets under the new-compound setting (a,d), the new-protein setting (b,e) and the both-new setting (c, f).

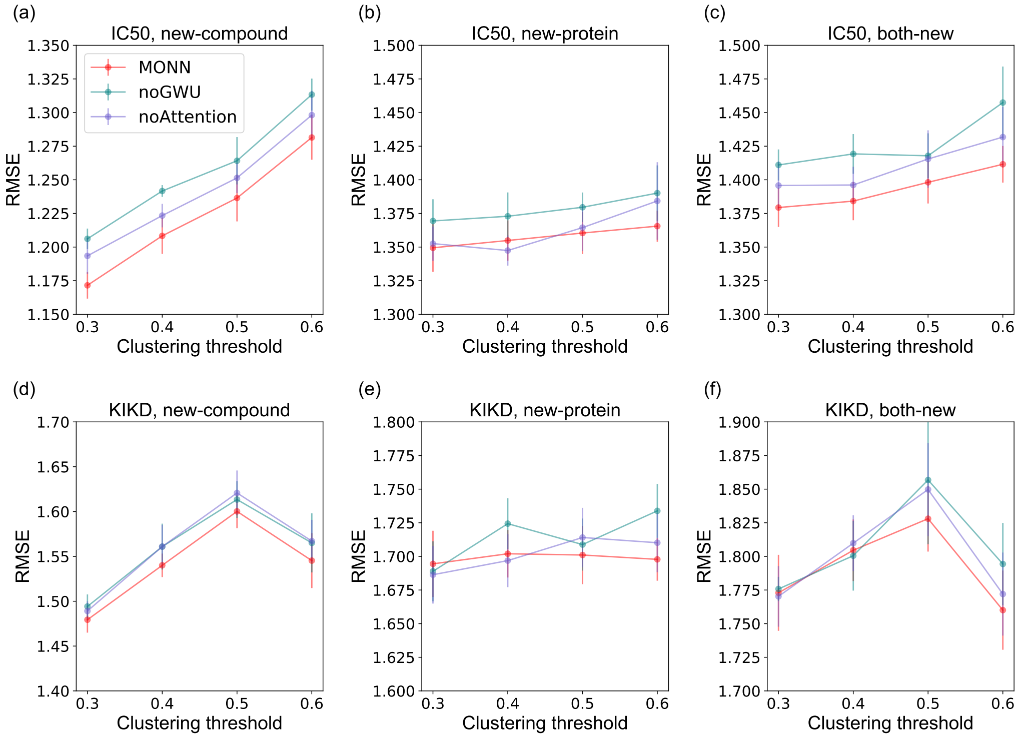


Figure S6: The ablation study for the binding affinity prediction task by removing the graph warp unit (GWU) or the attention part (DAN) of MONN. Performance was evaluated in terms of RMSE, using the IC50 (a-c) and KIKD (d-f) datasets under the new-compound setting (a,d), the new-protein setting (b,e) and the both-new setting (c, f).

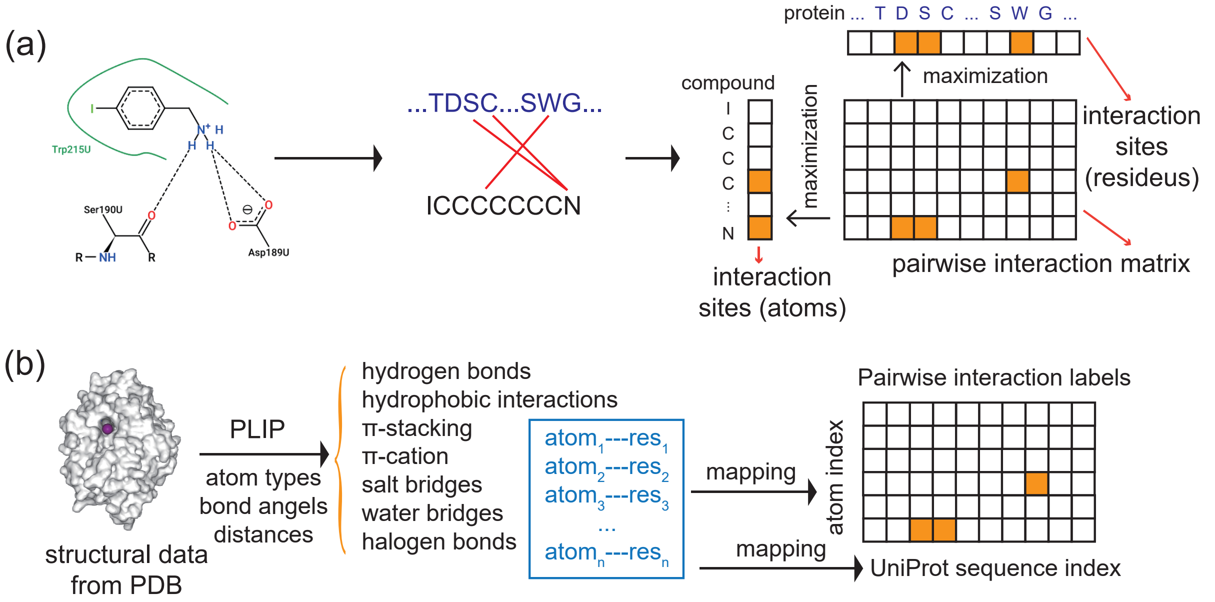


Figure S7: The schematic explanation for constructing pairwise interaction labels. (A) An example of pairwise interaction labels for a compound-protein complex (PDB ID: 5Z1C). The poseview figure is generated from https://poseview.zbh.uni-hamburg.de/5z1c. (B) The workflow for deriving the pairwise interaction labels from the structural data of a compound-protein complex.

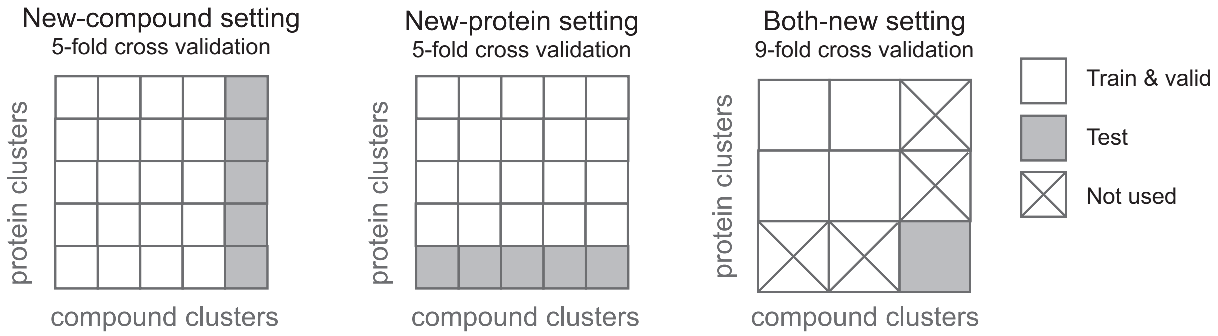


Figure S8: The three cross-validation settings.

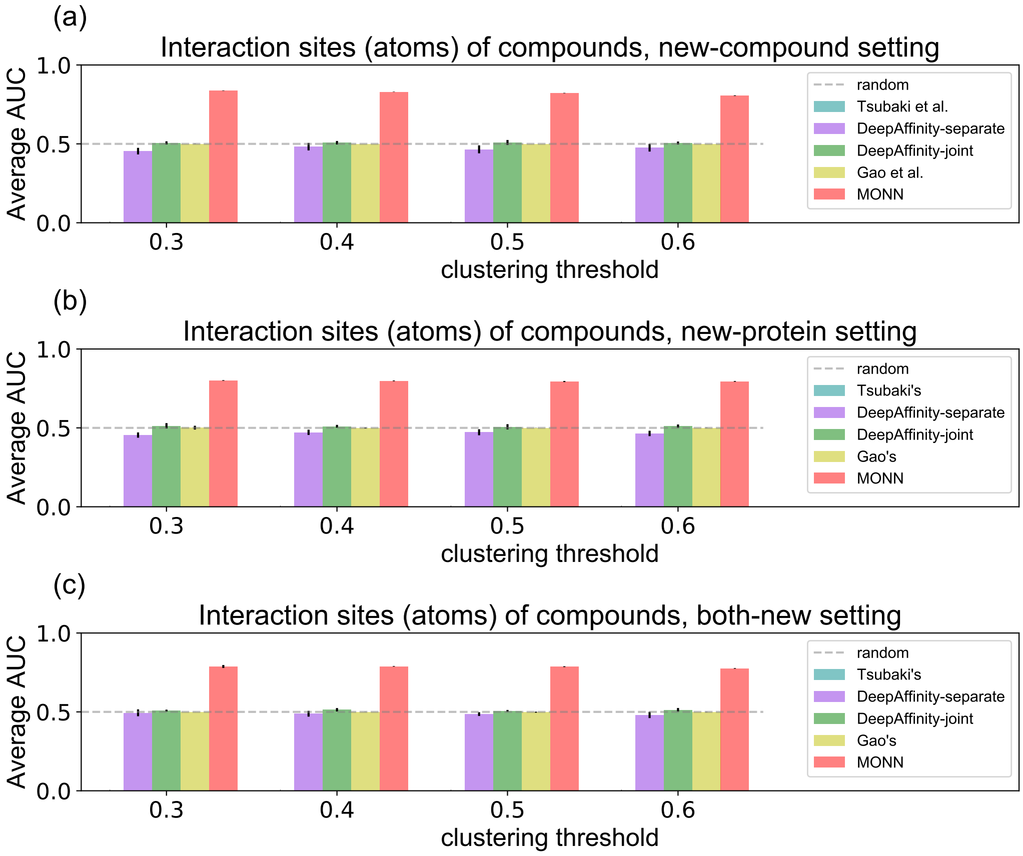


Figure S9: The average AUC scores for predicting interaction sites (atoms) of compounds using MONN and different neural attentions, under the new-compound setting (a), new-protein setting (b) and both-new setting (c), respectively. Note that Tsubaki et al.'s method is not applicable for this task and thus was not included in this figure.

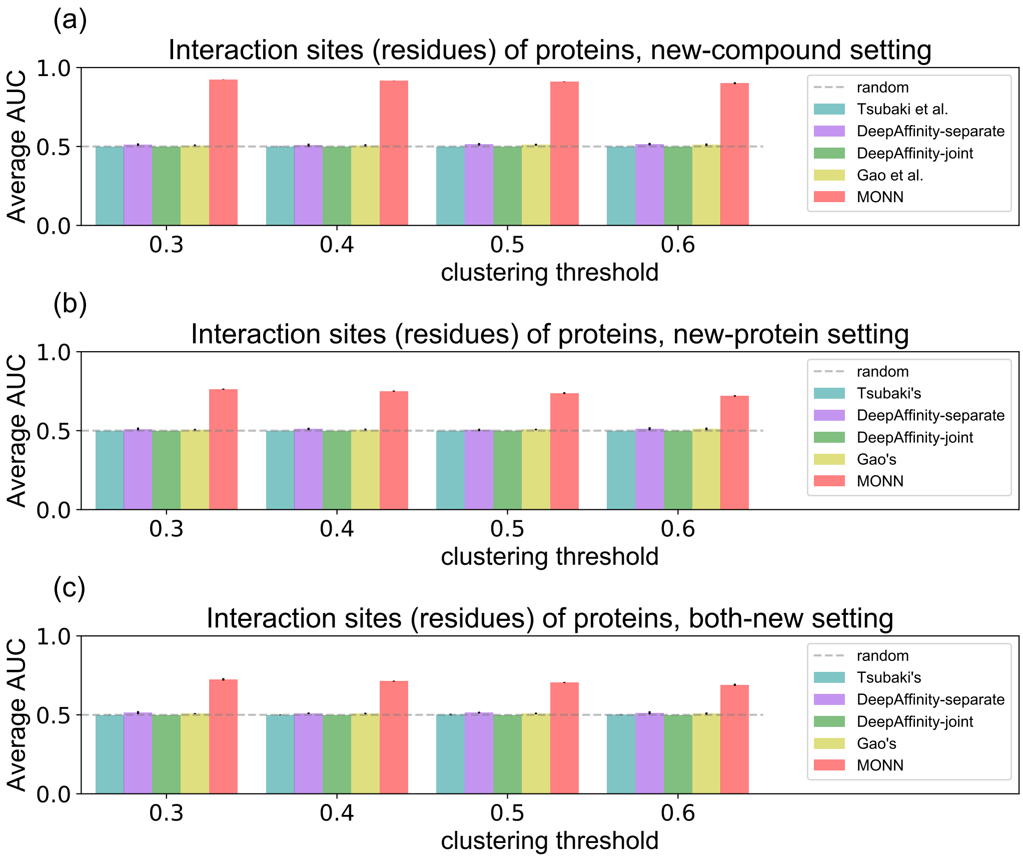


Figure S10: The average AUC scores for predicting interaction sites (residues) of proteins using MONN and different neural attentions, under the new-compound setting (a), new-protein setting (b) and both-new setting (c), respectively.

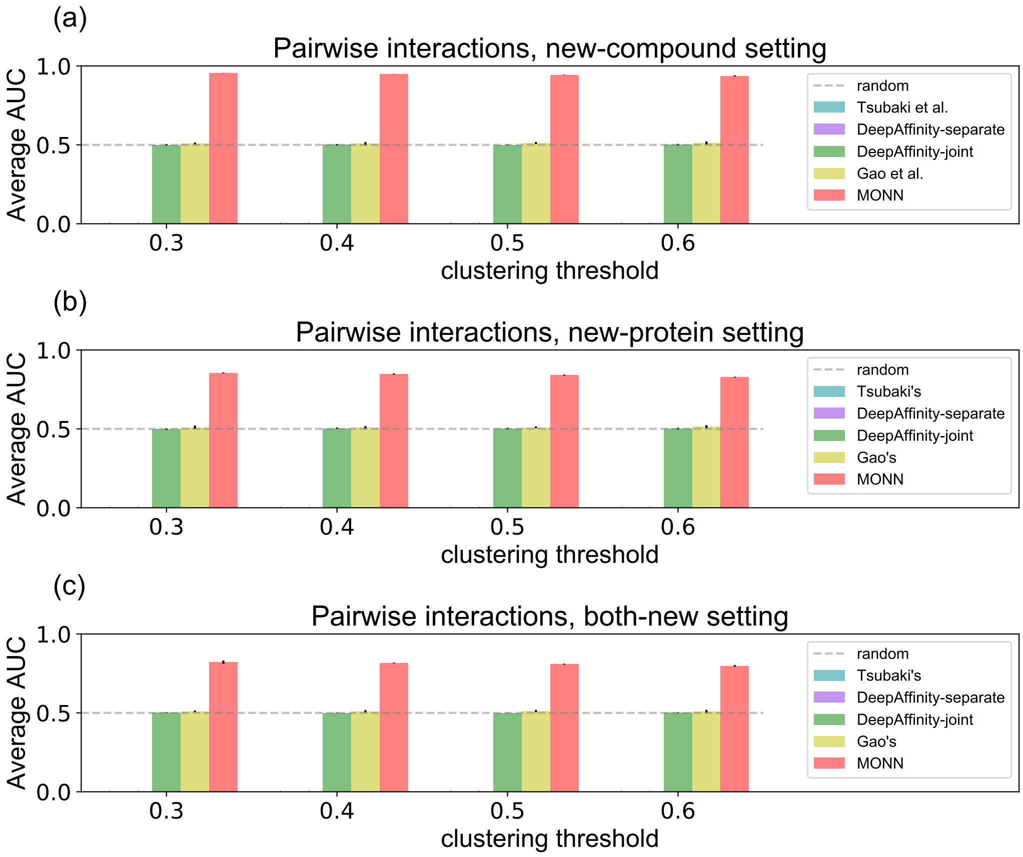


Figure S11: The average AUC scores for predicting pairwise interactions between compounds and proteins using MONN and different neural attentions, under the new-compound setting (a), new-protein setting (b) and both-new setting (c), respectively. Note that Tsubaki et al.'s method and DeepAffinity with separate attention are not applicable for this task and thus were not included in this figure.

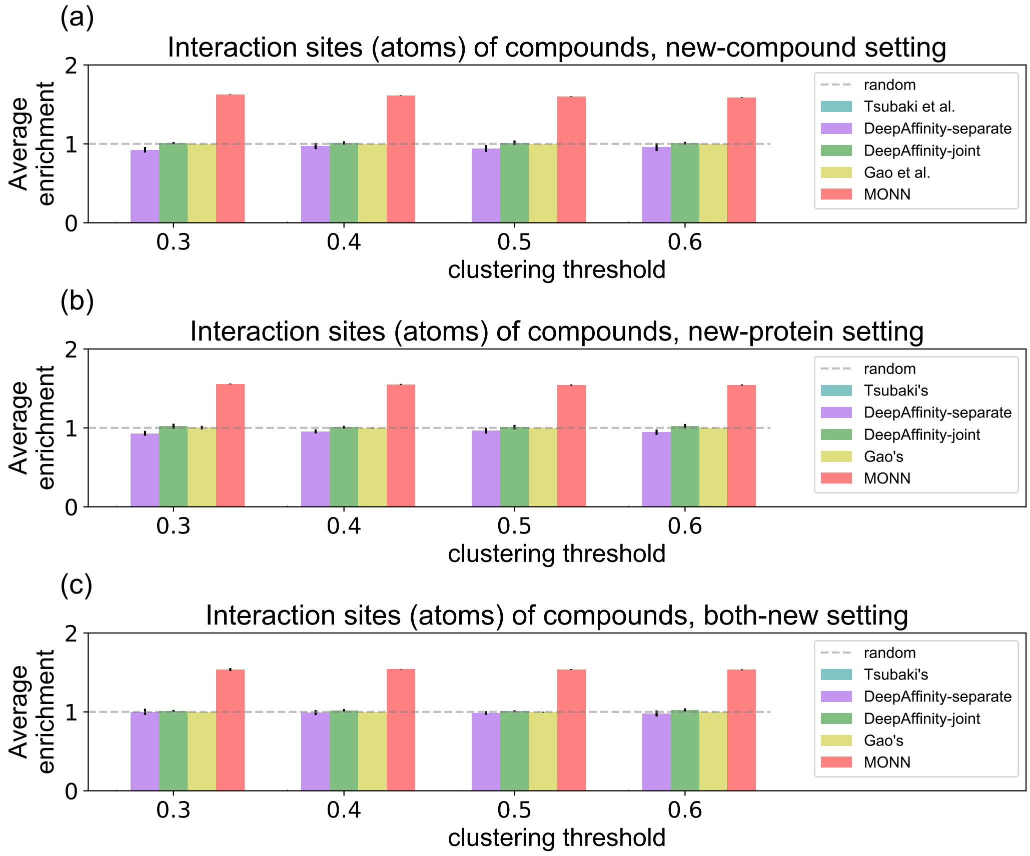


Figure S12: The average enrichment scores for predicting interaction sites (atoms) of compounds using MONN and different neural attentions, under the new-compound setting (a), new-protein setting (b) and both-new setting (c), respectively. Note that Tsubaki et al.'s method is not applicable for this task and thus was not included in this figure.

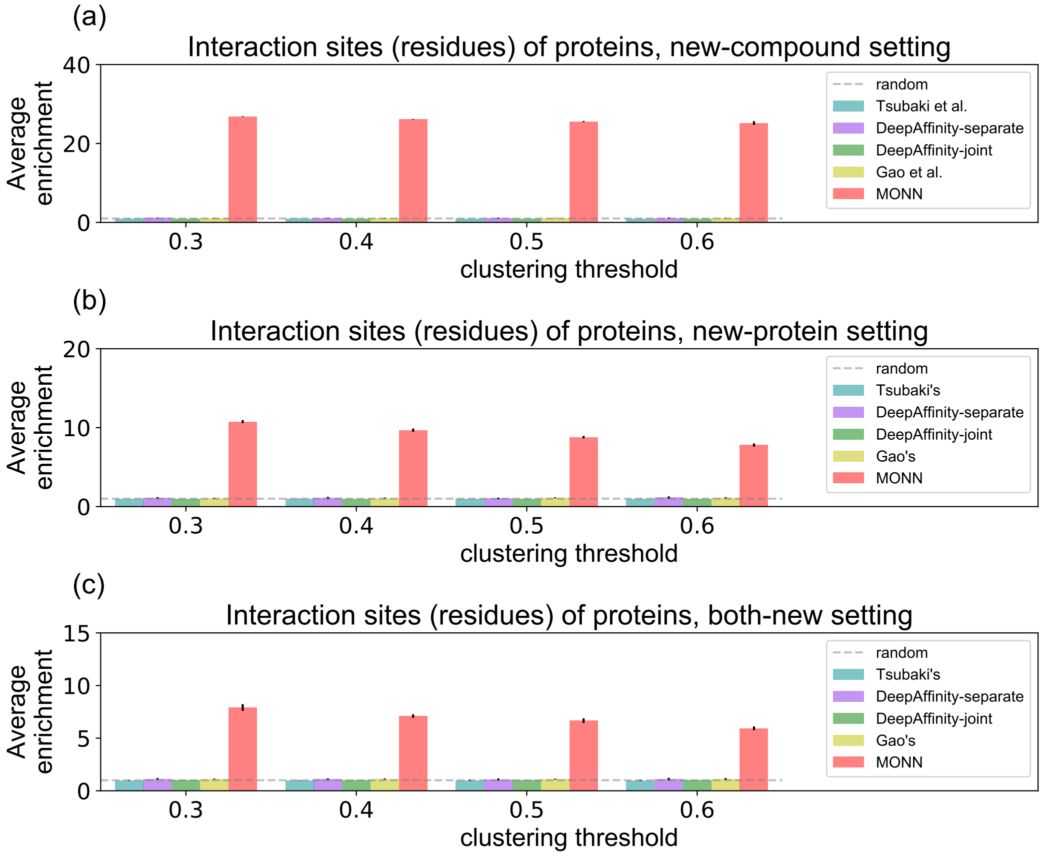


Figure S13: The average enrichment scores for predicting interaction sites (residues) in proteins using MONN and different neural attentions, under the new-compound setting (a), new-protein setting (b) and both-new setting (c), respectively.

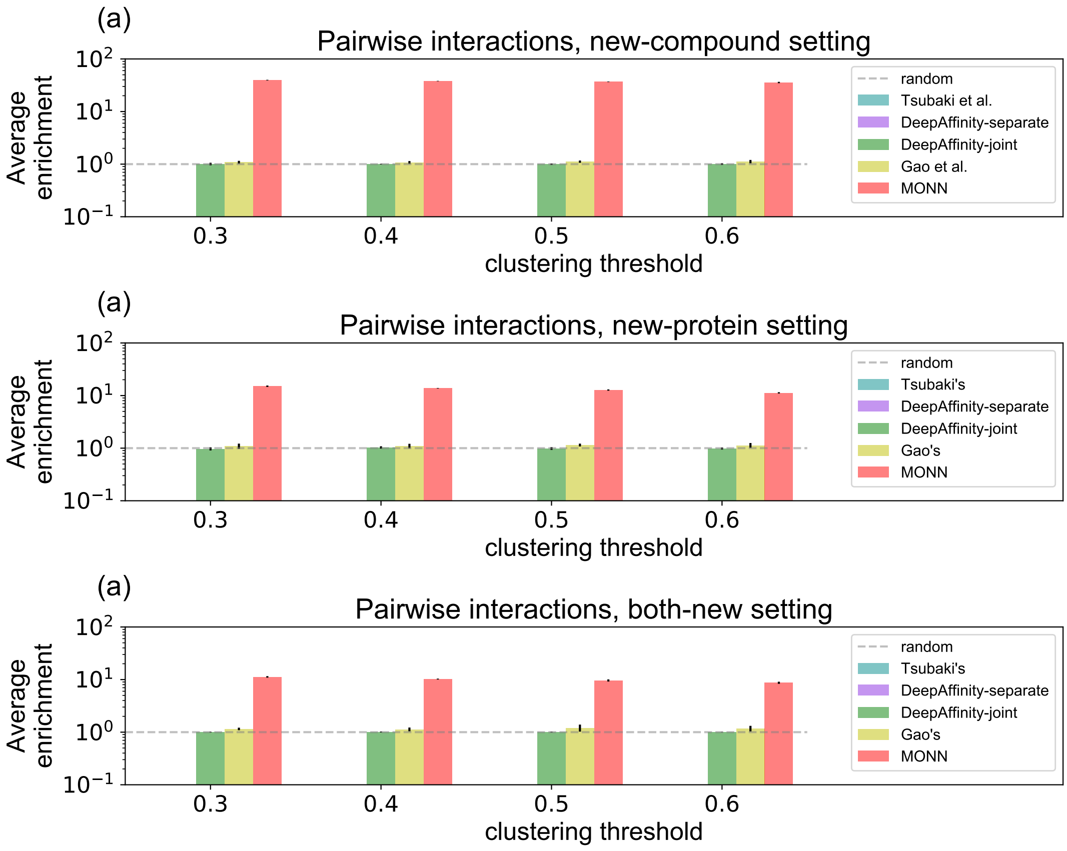


Figure S14: The average enrichment scores for predicting pairwise interactions between compounds and proteins using MONN and different neural attentions, under the new-compound setting (a), new-protein setting (b) and both-new setting (c), respectively. Note that Tsubaki et al.'s method and DeepAffinity with separate attention are not applicable for this task and thus were not included in this figure.

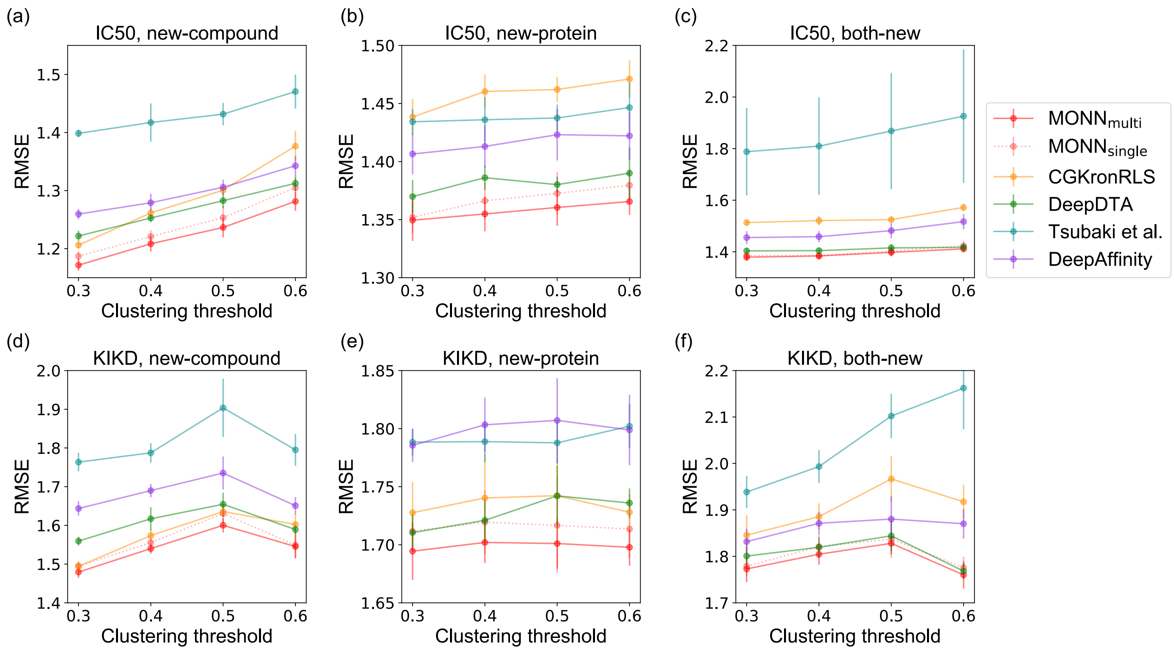


Figure S15: The performance of MONN and different baseline methods in predicting binding affinity values for the compound-protein pairs derived from the PDBbind, evaluated in terms of root mean squared error (RMSE). Cross-validation results for the IC50 (a-c) and KIKD (d-f) datasets under the new-compound setting (a,d), the new-protein setting (b,e) and the both-new setting (c, f) are shown.

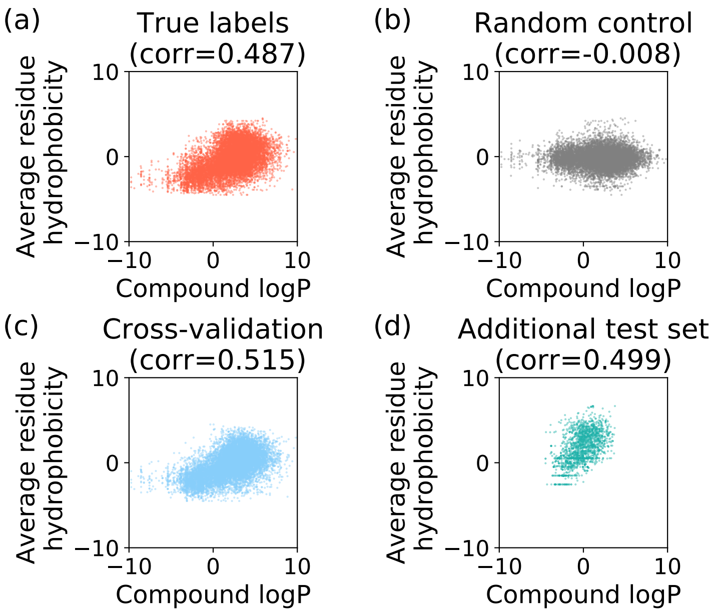


Figure S16: Correlation between the hydrophobicity scores of the compounds and the corresponding interaction sites in the proteins. (a) The interaction residues are derived from the pairwise interaction labels of the benchmark dataset. (b) The interaction residues of the proteins were derived from the randomly selected residues from the protein sequences. Here, the number of selected residues were the same as the number of true interaction sites in each protein sequence. (c) The interaction residues of the proteins were predicted by MONN. Using a nine-fold cross-validation on the benchmark dataset under the both-new setting with a clustering threshold 0.3, the interaction residues were derived from the predicted pairwise interaction matrices of the test samples for each fold. (d) The interaction residues were predicted for the compound-protein pairs in the additional test dataset, while the model was trained using the benchmark dataset.

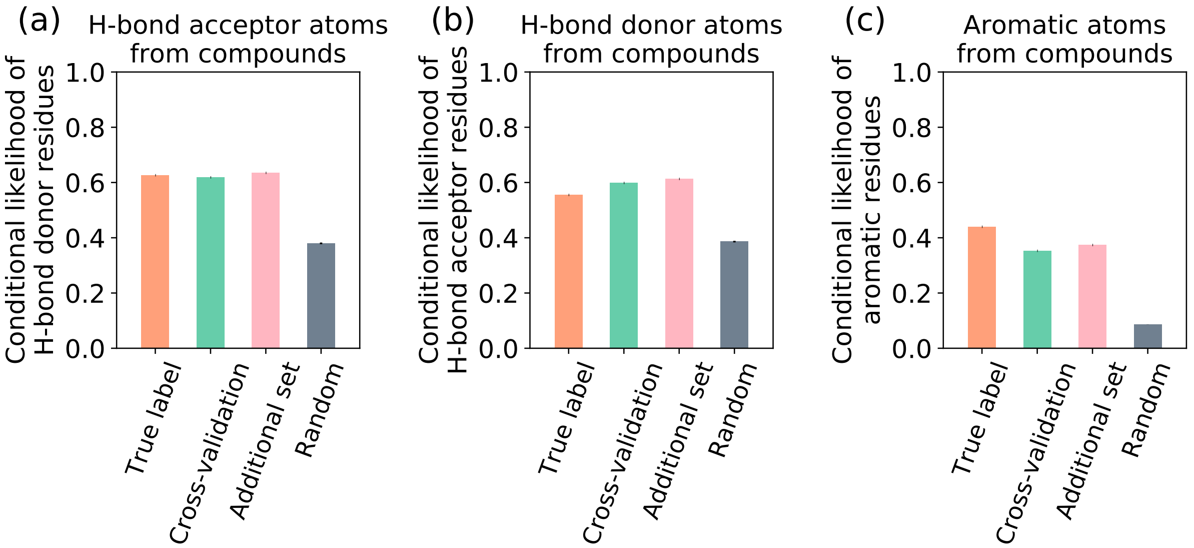


Figure S17: Conditional likelihood scores (whose definition can be found in the main text) measuring the preference of specific residue types given the property of interaction sites (atoms) of the compounds, including hydrogen-bond acceptor atoms (a), hydrogen-bond donor atoms (b) and aromatic atoms (c). Given a specific type of atoms from the compounds, we considered the interacting residues from four different situations, including true labels in the PDBbind dataset, the MONN predictions for all the test samples in a nine-fold cross-validation on the PDBbind dataset under the both-new setting with clustering threshold 0.3, the MONN predictions for the additional test dataset, and randomly selected residues along the same protein sequences.

[2] Katsuhiko Ishiguro, Shin Ichi Maeda, and Masanori Koyama. Graph warp module: an auxiliary module for boosting the power of graph neural networks. 2019.

[3] Junyoung Chung, Caglar Gulcehre, KyungHyun Cho, and Yoshua Bengio. Empirical evaluation of gated recurrent neural networks on sequence modeling. *arXiv preprint arXiv:1412.3555*, 2014.

[12] M Karimi, D Wu, Z Wang, and Y Shen. Deepaffinity: Interpretable deep learning of compound-protein affinity through unified recurrent and convolutional neural networks. *Bioinformatics (Oxford, England)*, 2019.

[13] Kyle Yingkai Gao, Achille Fokoue, Heng Luo, Arun Iyengar, Sanjoy Dey, and Ping Zhang. Interpretable drug target prediction using deep neural representation. In *IJCAI*, pages 3371–3377, 2018.

[14] Sebastian Salentin, Sven Schreiber, V Joachim Haupt, Melissa F Adasme, and Michael Schroeder. Plip: fully automated protein–ligand interaction profiler. *Nucleic acids research*, 43(W1):W443–W447, 2015.

[15] Renxiao Wang, Xueliang Fang, Yipin Lu, Chao-Yie Yang, and Shaomeng Wang. The pdbbind database: methodologies and updates. *Journal of medicinal chemistry*, 48(12):4111–4119, 2005.

[16] Renxiao Wang, Xueliang Fang, Yipin Lu, and Shaomeng Wang. The pdbbind database: Collection of binding affinities for protein- ligand complexes with known three-dimensional structures. *Journal of medicinal chemistry*, 47(12):2977–2980, 2004.

[17] Helen M Berman, John Westbrook, Zukang Feng, Gary Gilliland, Talapady N Bhat, Helge Weissig, Ilya N Shindyalov, and Philip E Bourne. The protein data bank. *Nucleic acids research*, 28(1):235–242, 2000.

[18] G Richard Bickerton, Gaia V Paolini, Jérémy Besnard, Sorel Muresan, and Andrew L Hopkins. Quantifying the chemical beauty of drugs. *Nature chemistry*, 4(2):90, 2012.
